# Inferring community architectures of multisensory pathways in *Drosophila* via unsupervised network embedding

**DOI:** 10.1101/2024.12.20.629573

**Authors:** Xiyang Sun, Fumiyasu Komaki

## Abstract

Understanding the complex architecture and functions of neural circuits is central to unraveling the mechanisms of multisensory integration. In this study, we analyzed the structural properties of the *Drosophila* adult brain to infer community structures within multisensory pathways. We adopt a network embedding method developed by ourselves, the Bidirectional Heterogeneous Graph Neural Network with Random Teleport (BHGNN-RT), designed to generate vector representations of neurons in a directed, heterogeneous brain connectome. This approach takes advantage of both structural connectivity and network heterogeneity features, enabling effective clustering of neurons and revealing hierarchical community architectures in olfactory and broader multisensory systems. We applied BHGNN-RT to the fly brain connectome to examine connectivity-based community organization in major neuronal classes along multisensory pathways, revealing distinct neural groups with unique connectivity patterns in the antennal lobe, lateral horn, mushroom body, and other brain regions. Further analysis showed how different neural groups contribute to the integration of sensory information in olfactory and multisensory systems. We also investigated the bilateral symmetry of the olfactory pathway, shedding light on how sensory signals are processed with ipsilateral and contralateral connections to ensure robust perception. Our findings demonstrate the utility of graph representation learning in analyzing the structural connectivity of complex neural systems. The insights gained from BHGNN-RT provide a deeper understanding of the community architecture in the *Drosophila* brain and contribute to a broader comprehension of the mechanisms underlying multisensory integration.

## Introduction

Understanding how the brain integrates sensory inputs from multiple sensory modalities to generate coherent perceptions and guide adaptive behaviors is one of the core challenges in neuroscience[1, 2]. Multisensory integration allows organisms to respond adaptively to complex environments by combining different types of sensory cues, such as visual [3], olfactory [4], mechanosensory [2], and other sensory cues [5]. Recent studies have demonstrated that multisensory integration enhances the accuracy and efficiency of perception by reducing ambiguities present in single-modality signals, thus supporting more robust decision-making and motor responses [6, 7, 8]. This process is fundamental for diverse tasks ranging from detecting predators to navigating toward food sources, making it a key area of study for understanding circuit organization and neural computation.

The fruit fly, *Drosophila melanogaster*, has served as a powerful animal model for studying the architecture and function of neural circuits that underlie multisensory integration [9, 10]. With its relatively small yet sophisticated nervous system [11, 12], *Drosophila* provides access to a complete connectome at synaptic resolution, enabling detailed mapping of neural pathways that integrate multiple sensory modalities [5]. In particular, the olfactory system of *Drosophila* has been extensively studied, revealing intricate details about interactions between olfactory sensory neurons (OSNs), antennal lobe neurons, and higher brain centers such as the mushroom body and the lateral horn [13, 14, 15]. These studies also emphasize the role of specific neurotransmitters, such as Acetylcholine (ACH) and Gamma-aminobutyric acid (GABA), in modulating sensory inputs and facilitating olfactory processing [16, 17]. Distinct sensory modalities typically interact or converge in specific brain regions, such as the antennal lobe, mushroom body, and central complex, to influence behavioral outcomes [5, 2]. The convergence of olfactory and gustatory inputs in the antennal lobe enables *Drosophila* to make more nuanced feeding decisions based on both smell and taste [4, 17]. Similarly, the interaction between the visual and olfactory pathways can enhance or modulate olfactory-driven behaviors [18]. Further, hygrosensory and thermosensory neurons, which respond to humidity and temperature, respectively, have been shown to interact with olfactory pathways to affect behaviors such as seeking optimal environmental conditions [5]. These studies indicate that the *Drosophila* brain uses sophisticated neural circuits to deal with multiple sensory modalities and produce behavioral outcomes, which are still being elucidated [5, 19].

Despite significant advances in connectome mapping and functional imaging, the intricate neural circuits about how neuronal groups form functional modules and how sensory modalities interact to form a cohesive sensory experience remain poorly understood [1, 20]. This is mainly due to the inherent complexity of biological networks, which are characterized by a large number of heterogeneous neurons, directed synaptic interactions, and non-uniform connectivity patterns [12, 21]. Traditional approaches, such as graph theory and morphological analysis, have revealed important aspects of brain organization but often fall short in capturing the heterogeneous nature and dynamics of neural connectivity, especially in multisensory systems where multiple modalities interact simultaneously [22, 1]. Recent studies have highlighted the importance of understanding both the structural and functional aspects of these neural circuits to gain a complete picture of sensory integration [23, 8, 24]. This twofold perspective not only enhances our comprehension of basic sensory functions but also informs the development of interventions for sensory-related disorders. Hence, it highlighted the requirements of advanced computational tools that can account for the directionality and diversity of synaptic connections, as well as the unique features of biological networks.

Graph Neural Networks (GNNs) have recently gained prominence as a powerful tool for modeling complex graph-structured data in many scenarios, including biology, social networks, and chemistry [25, 26, 27]. The fundamental concept behind GNNs is to extend traditional neural networks to graph data, allowing each node to aggregate information from its neighbors through iterative message-passing mechanisms [28, 29]. This iterative nature of message-passing enables GNNs to capture both local and higher-order topological features, making them particularly well-suited for analyzing connectome-like data and extracting meaningful insights about neural connectivity. Recent advancements have led to diverse GNN variants, such as Graph Convolutional Network (GCN) [30], Graph Attention Network (GAT) [31], GraphSAGE [32], each contributing unique capabilities to improve feature extraction, scalability, and generalizability in large and complex graphs [33]. Some of them have been successfully applied in network neuroscience, such as missing brain graph synthesis, disease classification, and the integration of population graphs [34, 35]. However, most existing GNN-based methods struggle with challenges unique to biological networks, particularly in handling directed and heterogeneous connections that characterize synaptic interactions in the brain [33, 36]. There is a pressing need for specialized GNN models that can effectively capture these properties in a way that preserves biologically relevant features.

To address these challenges, we adopt the recently proposed Bidirectional Heterogeneous Graph Neural Network with random Teleport (BHGNN-RT), an approach introduced by ourselves [37] for graph embeddings of general directed graphs, which aligns well with the unique needs of biological neural circuits. BHGNN-RT employs a bidirectional message-passing strategy to account for the different roles of incoming and outgoing signals, which is crucial for preserving the directionality of information flow in neural circuits. Additionally, the model includes an attention mechanism to weigh the significance of network heterogeneity and a random teleport component to mitigate the over-smoothing issue common in deep graph models. We applied BHGNN-RT to the *Drosophila* brain connectome to reveal the hierarchical structure and community organization within key sensory regions, known for their roles in multisensory integration. By analyzing these community architectures, we aimed to understand how different sensory modalities converge to influence behavior. This research not only enhances our knowledge of *Drosophila* neural circuits but also offers a generalizable framework for analyzing biological networks that may inform future studies in neuroscience and artificial intelligence.

## Results

### Graph representation learning for biological neural networks

Considering the complexity of the structural connectivity in the biological neural circuits, how can we efficiently represent neurons with diverse biological properties and infer the overall hierarchical architecture? Our analytical strategy is to adopt a novel network embedding method to capture the intricate properties of the neural circuits. This method generates informative vector representations for neurons that incorporate local and global connectivity patterns, which is vital for understanding the complex dynamics within these circuits. It enables the identification of fine-grained community structures and provides insight into the hierarchical organization of the brain connectome (Fig.1A).

**Figure 1.**
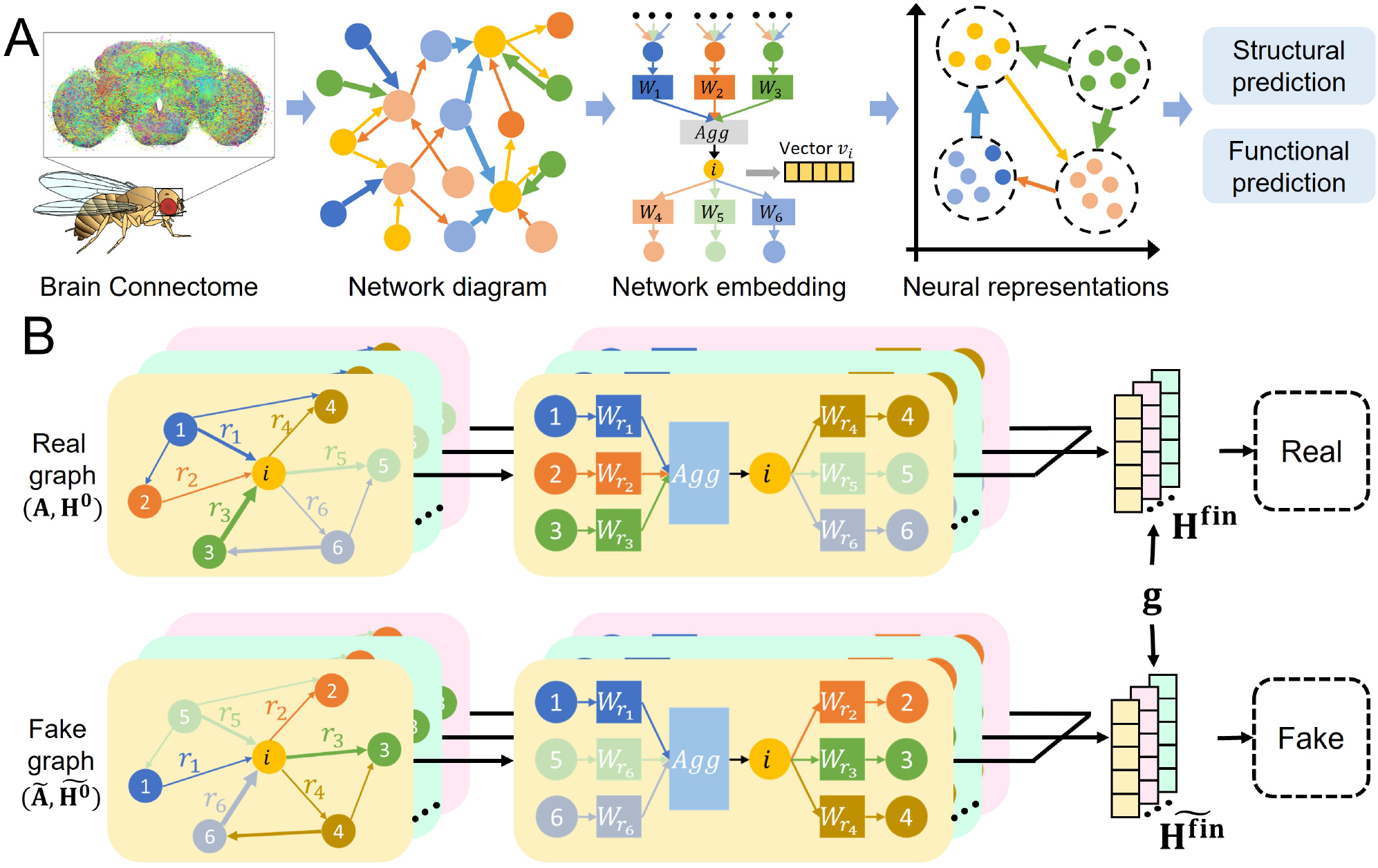
Network representation learning based on neuronal connectivity patterns. (A) A schematic of our analytic strategy based on network embedding. Considering the *Drosophila* brain connectome as a directed, weighted, and heterogeneous graph, the network embedding strategy is utilized to learn the neural vector representation. Based on these neural representations, we can infer the structural and functional roles of the neuron community groups. (B) The overall framework of BHGNN-RT for graph representation learning. In the directed, weighted, and heterogeneous networks, an example neuron *i* is linked to neighboring neurons under different edge relations (*r*_*i*_). The neural representation is updated based on its connectivity pattern and network heterogeneity. Each edge for message-passing is weighted by a matrix 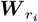 for different edge relations. The objective function in BHGNN-RT is to optimize the mutual information between neural representation *h*_*i*_ and the graph summary *g*.

#### Task definition: Encoding vector representations for neurons

To analyze the *Drosophila* brain connectome (FlyWire dataset), we first formulate it as a directed, weighted, and heterogeneous graph 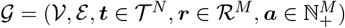, where each node represents a neuron and each edge represents a chemical synapse. The node set 𝒱 consists of *N* = 134, 191 neurons, with each node labeled as *v*_*i*_ for *i* = 1, …, *N*. Here, *t*_*i*_ ∈ 𝒯, the *i*-th element of ***t***, denotes the neural type. The edge set ℰ contains *M* = 2, 831, 350 directed edges of the form (*v*_*i*_, *v*_*j*_). Each edge (*v*_*i*_, *v*_*j*_) in ℰ is assigned a unique index [*i, j*] ∈ {1, …, *M*}, where the edges are ordered lexicographically; that is, *i* takes precedence over *j* in the ordering. The synaptic weight *a*_[*i*,*j*]_ ∈ ℕ_+_, the [*i, j*]-th element of ***a***, represents the number of synapses in the connection (*v*_*i*_, *v*_*j*_). The edge type *r*_[*i*,*j*]_ ∈ ℛ, the [*i, j*]-th element of ***r***, characterizes the synaptic relationship and is defined by distinct neurotransmitters. The set 𝒯 consists of 7 distinct neural types, and the set ℛ consists of 6 types of synaptic relations. The adjacency matrix ***A*** ∈ ℝ^*N* ×*N*^ represents the neural connectivity, with entries defined as

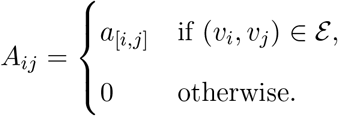

The objective here is to construct numerical vectors corresponding to each node of the graph. These vectors effectively capture and preserve the role of each node in the graph. Although graphs, in their original form, are not straightforward to handle numerically or statistically, their conversion into vectors enables various forms of analysis. The assignment of numerical vectors to all nodes is called network embedding or graph embedding, which has recently attracted significant attention in machine learning and artificial intelligence. For network embedding, we construct an unsupervised encoder, *Enc*(***A, H***^0^) = ***H***^fin^, where 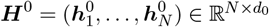 represents the initial neural representations with a predefined dimension *d*_0_, initialized as all-ones vectors. The output, 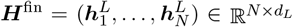, is the final learned neural representations. Here, *L* denotes the number of iterations, as described in the next subsection.

#### Bidirectional Heterogeneous Graph Neural Network with Random Teleport (BHGNN-RT)

Inspired by the network characteristics (Supplementary Fig.1), we applied the node-embedding algorithm, Bidirectional Heterogeneous Graph Neural Network with Random Teleport (BHGNN-RT), proposed for general directed, heterogeneous graphs. This approach captures the bidirectional message-passing process and network heterogeneity, which is essential for this directed, weighted, and heterogeneous brain connectome. An overview of our approach based on BHGNN-RT is provided below in this section. For details on BHGNN-RT, see [37].

For each neuron *v*_*i*_, BHGNN-RT distinguishes between its upstream neurons *N*_in_(*v*_*i*_), which is the set of nodes with directed edges pointing to *v*_*i*_, and downstream neurons *N*_out_(*v*_*i*_), which is the set of nodes that *v*_*i*_ points directed edges to (Fig.1B). This distinction is particularly important in the context of biological networks, where afferent and efferent pathways carry different types of messages [38, 39]. Additionally, this model incorporates heterogeneous node and edge types, accounting for the diversity of neuron types and synaptic interactions in the connectome.

The vector update procedure described below is repeated *L* times. In this study, we set *L* = 4. In the field of graph neural networks, the *l*-th iteration is called the *l*-th layer. After *L* layers of iteration, the final neural embedding is obtained as 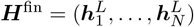.

In each layer, for each neuron *v*_*i*_, the contributions from its upstream neurons *N*_in_(*v*_*i*_) are aggregated to update its representation. Specifically, the aggregated input is calculated as

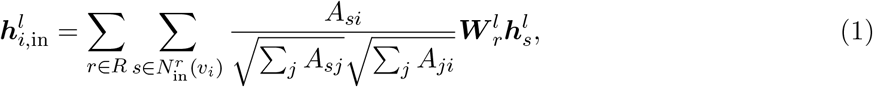

where 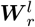 is a *d*^*l*+1^ × *d*^*l*^ weight matrix associated with the synaptic type *r* (Fig.1B), which is optimized during the learning process, and 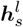 is the representation of neuron *v*_*s*_ at layer *l*. Here, the *d*^*l*+1^-dimensional vector 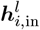 represents the aggregated contributions from upstream neurons to the representation of *v*_*i*_ at the *l*-th layer.

Similarly, the contributions to downstream neurons from neuron *v*_*i*_ are aggregated as

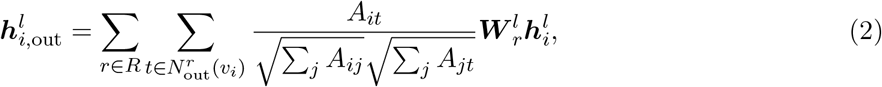

where 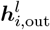 represents the aggregated contributions from *v*_*i*_ to its downstream neurons at the *l*-th layer.

The weight matrix 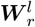 is the same as that used in Eq.1.

After calculating 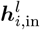 and 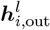, they are aggregated to update the neural representation. The unnormalized vector representation of neuron *v*_*i*_ at layer *l* + 1 is

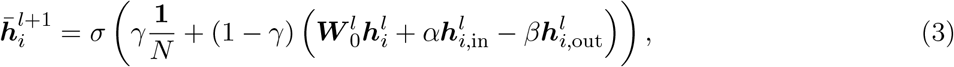

where the activation function *σ* is a parametric rectified linear unit (PReLU), defined as *σ*(·) = max(0, ·) + *k* min(0, ·) with a learnable parameter *k*, and 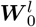 is a *d*^*l*+1^ × *d*^*l*^ weight matrix optimized during the learning process. The hyperparameters *α, β*, and *γ* ∈ [0, 1] control the contributions of different message components and are optimized during training. The term *γ***1***/N* is introduced to ensure the stability of the estimated vector. The updated neuron embedding is then normalized as 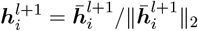, where 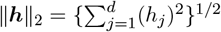 is the standard Euclidean norm.

#### Objective function

To achieve unsupervised learning, we adopt an objective function inspired by deep graph infomax (DGI), which uses a quantity motivated by mutual information (MI) between the neural representations and a network summary ***g*** defined below [40]. In parallel with the FlyWire connectome 𝒢, we construct a fake graph. Let *τ* (*i*) (*i* = 1, …, *N*) be a random permutation of *i* = 1, …, *N*. The fake graph is defined as 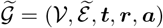, where the edge set 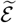 consists of edges (*v*_*τ* (*i*)_, *v*_*j*_) for (*v*_*i*_, *v*_*j*_) ∈ ℰ. We use an initial feature matrix 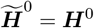.

Real output of BHGNN-RT for the real graph 𝒢 is ***H*** ∈ ℝ^*N* ×*d*^, and the fake output is 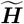. The network representation ***g*** ∈ ℝ^*d*^ for 𝒢 is derived as 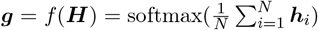 by averaging all neural representations in the brain connectome.

As a proxy for maximizing the mutual information between neuron-network pairwise representations, a discriminator function 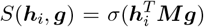 is applied to calculate the probabilistic score for the neuron-network pair, where 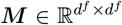 is a learnable scoring matrix and *σ* is a sigmoid function. Concerning the set of original and fake graphs, the log-likelihood function is configured as

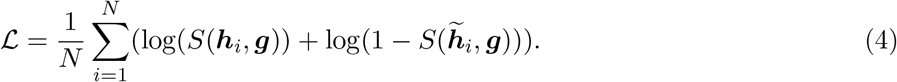

This logarithmic likelihood value is motivated by mutual information and assigns higher scores to positive embeddings and lower scores to negative ones. The objective function encourages the encoder to capture meaningful information shared between all neurons (Fig.1B).

The performance of BHGNN-RT was verified in benchmark data sets, while compared with other state-of-the-art algorithms (details in Materials and Methods). In the task of unsupervised clustering, the results demonstrate the versatility and effectiveness of BHGNN-RT in handling complex heterogeneous networks, capturing both structural and nodal information more effectively compared to other methods [37]. BHGNN-RT helps generate nodal clusters with clearer clustering boundaries and a higher SIL score. Hence, our method is promising to improve unsupervised clustering quality while capturing comprehensive nodal connectivity profiles and graph-level structural properties.

### Connectivity-based community inference of Kenyon Cells

We apply BHGNN-RT on Kenyon Cells to examine its potential for connectivity-based clustering of neural groups in biological neural circuits. The superior performance of BHGNN-RT on benchmark datasets has been verified in [37]. Kenyon Cells (KCs), the intrinsic neurons in the mushroom body (MB), are selected mainly because of their well-defined spatial locations and structural organization in MB neuropils [41, 42, 12]. However, the complexity within the connectivity patterns of the KCs makes it difficult to uncover their functional roles within the MB [43]. Therefore, it is highly significant to understand the organizational principles of KCs based on connectivity-based analysis to uncover the neural mechanisms underlying adaptive behaviors of *Drosophila*.

For *Drosophila* adult, 5177 KCs have been revealed in its brain connectome, with 2,597 KCs in the right hemisphere and 2,580 KCs in the left (Fig.2A) [11]. Based on the vector representations via BHGNN-RT, we inferred community structures and clustered KCs into 11 distinct neural groups, as shown in Fig.2B-D. The optimal number of clusters was determined using the elbow method, where the distortion score indicated that 11 clusters captured the complexity of the KC population most effectively (Fig.2B). We aligned the hierarchical clustering dendrogram with the compartmentalization of the MB lobes (*α, β, γ*), according to the previous annotations [44, 45]. Corresponding spatial arrangements of these 11 KC clusters are depicted in Fig.2E. Most neural clusters exhibit a distribution with 40∼60% neurons localized to unilateral brain regions, indicating a relatively symmetric organization across two hemispheres (Fig.2F). Overall, this connectivity-based clustering can distinguish neuronal groups with distinct neighboring neurons even if they are in different hemispheres or located in the same MB lobes with similar morphologies.

**Figure 2.**
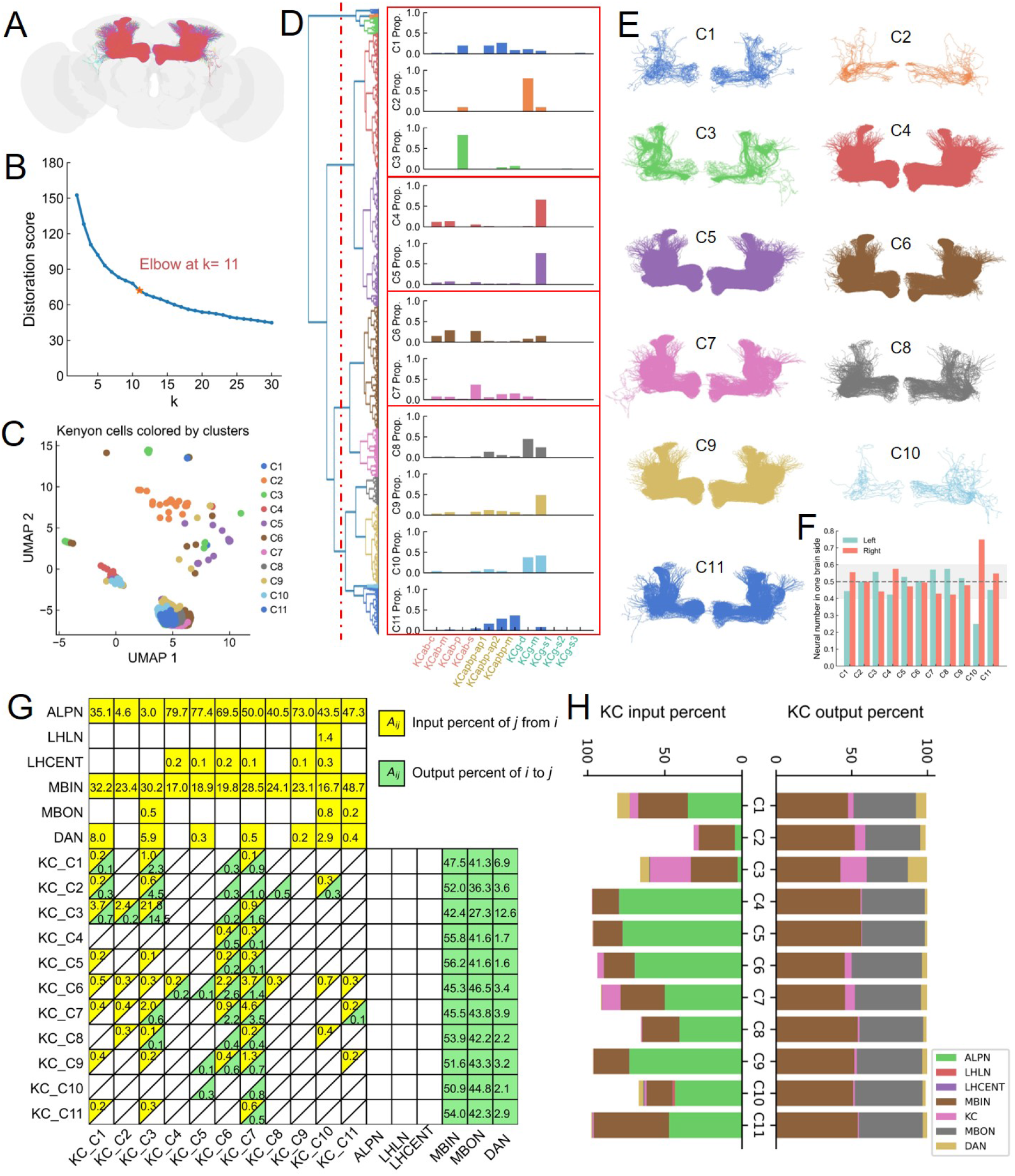
Investigation of hierarchical clustering results based on Kenyon Cells (KCs). (A) The spatial organization of KCs in Mushroom Body (MB). (B) The elbow method is utilized to determine the coarseness of clusters, based on vector representations of KCs via BHGNN-RT. (C) The UMAP visualization of vector representation for KCs, colored by the cluster labels. (D) Hierarchical clustering of KCs based on their vector representation (Left panel), and their distributions of cell type based on the ground truth. (E) Spatial organizations of individual clusters of KCs. (F) Neuronal number of each KC cluster in the hemisphere. (G-H) The percentage of synaptic input and output of KC clusters, when considering its connectivity with the major neuronal classes along the olfactory pathway. (G) displays the detailed value of input and output percentage, which is larger than 0.1%.

Within the clustering results, seven KC clusters (C2, C3, C4, C5, C8, C9, and C10) display dominant KCab-p, or KCg-d or KCg-m neural types, which are also the primary cell types of KCs annotated in [45]. However, the connectivity patterns of these clusters vary. The cluster C3, mainly for KCab-p neurons, exhibits a strong preference for recurrent connections (21.5% neural input) within itself (Fig.2G-H). This intracluster connectivity pattern is supported by the concentration of the neurite arborizations of these KCab-p neurons in posterior *α, β* lobes, which suggests a role in specialized output modulation [46, 47]. Besides, both clusters C2 and C3 only receive a small fraction of inputs (<5%) from the antennal lobe projection neurons (ALPNs), much smaller than other KC clusters. Clusters C4 and C5 exhibit high similarity in their neural components, where KCg-m neurons are dominant. As one of the primary recipients of olfactory input, these KCg-m neural groups (C4, C5, C9) gained at least 70% input from ALPNs (Fig.2G).

In Fig.2G-H, the distinct connectivity profiles of these KC clusters suggest that different neural groups may be specialized to perform diverse roles in sensory information processing and associative learning. Here, we propose some relevant hypotheses for their functional roles in the *Drosophila* brain connectome. First, neural clusters C6 and C7, mainly for KCab and KCapbp neurons, present widespread connections to other KC clusters, with both incoming and outgoing connections. This may suggest that these neurons specialize in recurrent connections within KCs and associative learning. Secondly, neural clusters C2, C8, and C10, as the mixture of KCg-d and KCg-m neurons, may serve distinct roles in sensory information processing. Cluster C2 receives little input from ALPNs, but Clusters C8 and C10 still receive around 50% incoming messages from ALPNs. Scheffer’s work stated that KCg-d neurons primarily receive optic input, while other KCg neurons get primarily olfactory input [44, 48]. Hence, we suppose cluster C2 is more inclined to deal with non-olfactory inputs while clusters C8 and C10 may serve as integrating multisensory information.

Furthermore, we compared this connectivity-based clustering result with morphology-based clustering via NBLAST [22]. Supplementary Fig.3 presents the hierarchical clustering of KCs solely based on neural morphological similarity. Based on the FlyWire annotations, the distribution of cell types within these six neuronal groups can be simply categorized into three main classes, including KCab, KCapbp, and KCg (Supplementary Fig.2B). Only the third cluster covers both KCab and KCapbp neurons. These neuronal clusters typically follow their spatial localizations in different lobes (*α, β, γ* lobes) of MB. However, this morphology-based clustering cannot tell the difference in connectivity profiles of these neurons, which are important insights for infer neural functions. What’s more, NBLAST treats neurons in the left and right hemispheres completely different, while our embedding method can still consider the similarity of neural connectivities within both hemispheres.

Our connectivity-based clustering and analyses reveal a highly structured and heterogeneous organization of KCs within the Mushroom Body. The identification of 11 distinct clusters, each with unique cell type compositions and connectivity patterns, highlights the complexity and specialization of KCs. These analyses provide valuable insights into the neural connectivity and functional organization of the MB, contributing to a deeper understanding of the neural basis of cognitive processes in the brain.

### Community architectures across neuronal classes in olfactory pathway

To extend our analysis beyond KCs, we further investigated the neural community architecture along the olfactory pathway and how they contribute to olfactory processing and integration in the *Drosophila* brain. In particular, we focus on the major neural classes, including olfactory sensory neurons, antennal lobe local neurons (ALLNs), antennal lobe projection neurons (ALPNs), lateral horn local neurons (LHLNs), lateral horn centrifugal neurons (LHCENTs), mushroom body input neurons (MBINs), Kenyon cells (KCs), mushroom body output neurons (MBONs), dopaminergic neurons (DANs), and central complex neurons (CX). Although significant progress has been made in understanding the olfactory pathway in the *Drosophila* adult brain [41, 14, 12], several limitations persist in current studies. One major limitation, the incomplete mapping of synaptic connections at the cellular level, has now been overcome by the FlyWire dataset, which leverages comprehensive electron microscopy [11]. Another limitation involves the functional characterization of neurons. While anatomical studies provide extensive data on the structure and connectivity of olfactory neurons, functional studies often struggle to correlate these structural data with precise functional roles due to the complexity and variability of olfactory stimuli [12, 49]. These gaps hinder the ability to fully understand how specific neural circuits translate neural signals layer by layer.

The synaptic connections across these ten neural classes are depicted along the fly olfactory pathway (Fig.3A), involving a highly organized flow of information from olfactory sensory neurons to higher brain regions. Starting from the antennal lobe (AL), the olfactory sensory neurons directly synapse with the local neurons (ALLNs) and projection neurons (ALPNs). Although the majority of olfactory output (∼10^5^ synapses) is to ALPNs, these sensory neurons receive few direct reciprocal connections (∼10^2^ synapses) from ALPNs (Fig.3A, Fig.4A). Within the AL, the mutual interactions (∼10^5^ synapses) between ALLNs and ALPNs remain the strongest (Fig.3A, Fig.4B-C). Sensory information is then relayed from ALPNs to neurons in the lateral horn and mushroom body, while the majority of ALPN output(∼10^5^ synapses, especially from ALPN cluster C3) is to KCs through a complex synaptic network in the calyx of MB (Fig.3A, Fig.4C) [43]. Within the MB, the strongest reciprocal connections (∼10^5^ synapses) occur between MBINs and KCs (Fig.3A, Fig.4F). The processed information is then conveyed from KCs to MBONs (∼10^5^ synapses), with little feedback (∼10^2^ synapses) from MBONs (Fig.3A, Fig.4G). Outside the mushroom body, KCs project much information to DANs (>10^4^ synapses). For the lateral horn, the LHLNs and LHCENTs present a not very strong mutual interaction (∼10^3^ synapses) between each other (Fig.3A). However, LHLNs provide heavy feedback (∼10^4^ synapses) to ALPNs, much larger than other neural classes in the lateral horn and mushroom body (Fig.3A). Further, sensory information is delivered to higher brain regions (such as the central complex, etc) from ALPNs, LHCENTs, MBONs, and DNAs. Among these neural classes along the olfactory pathway, MBINs present the highest degree and eigenvector centralities (Fig.3C-D), indicating their significant importance in aggregating and relaying diverse sensory information to the mushroom body.

**Figure 3.**
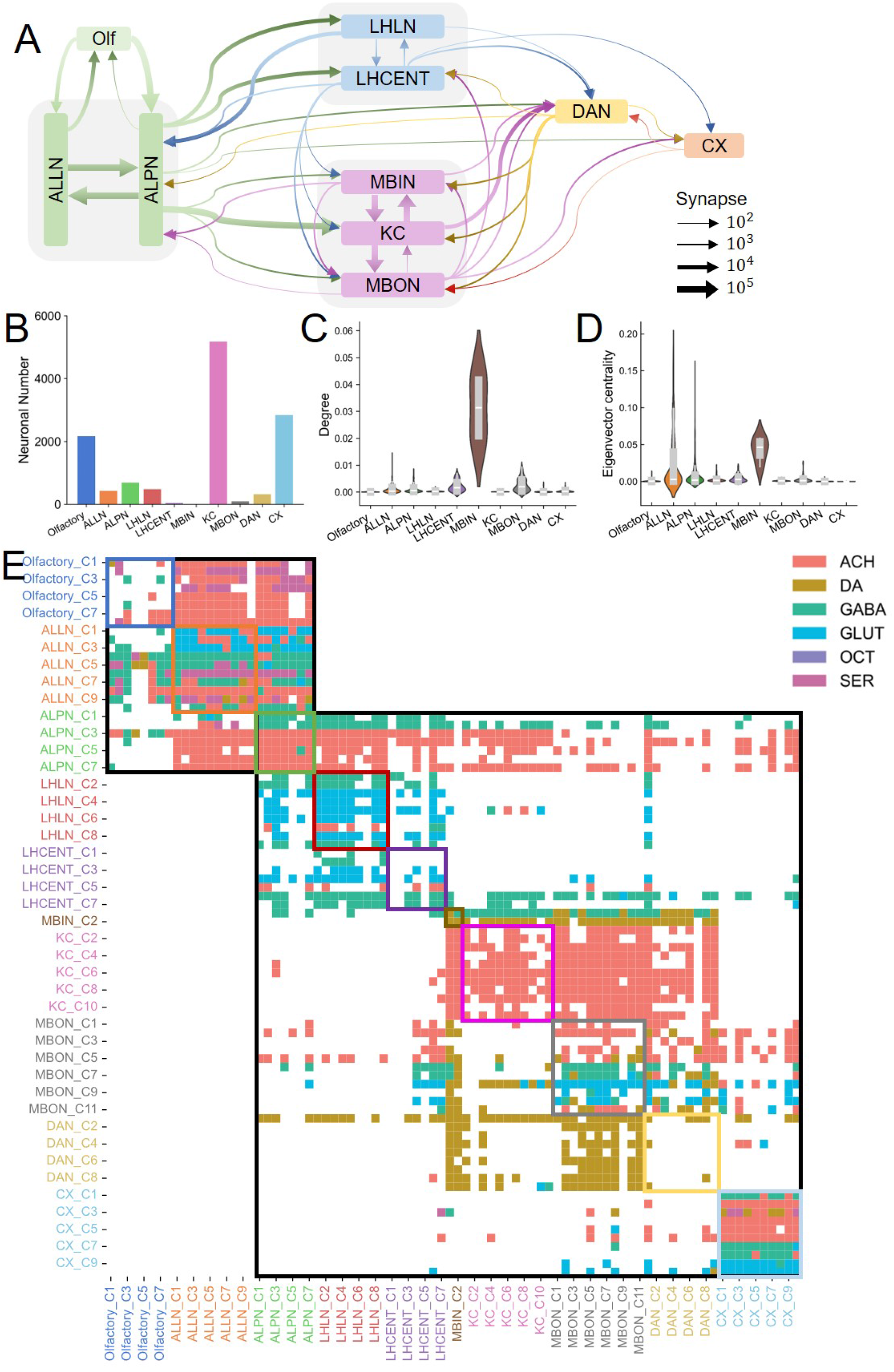
Analysis of community architecture along the olfactory pathway. (A) A diagram for the synaptic connectivities within the main neuronal classes along olfactory pathway, including olfactory neuron, Antennal lobe local neuron (ALLN), Antennal lobe projection neuron (ALPN), Lateral horn local neuron (LHLN), Lateral horn centrifugal neurons (LHCENT), Mushroom body input neuron (MBIN), Kenyon cell (KC), Mushroom body output neuron (MBON), Dopaminergic neuron (DAN) and central complex neuron (CX). Only the interclass synaptic connections with a strength larger than 100 are displayed here. (B) Neuronal number of these major neuronal classes. (C) Centrality organizations of these main neuronal classes, including degree centrality and eigenvector centrality. (D) The connectivity matrix of clusters of the above neuronal classes. The cluster results are inferred based on neural vector representations via BHGNN-RT, following similar procedures in Fig.3B-D. Each element in the connectivity matrix is the dominant synaptic type between two neural clusters.

Based on the neural vector representations via BHGNN-RT, we generated the clustering results for these ten major neural classes (Supplementary Fig.4). Fig.3E describes the connectivity matrix of the corresponding neural clusters, whose element represents the dominant synaptic type between pairwise neural groups. This organization pattern of the connectivity matrix characterizes a typical bow-tie architecture along the olfactory pathway. In such a bow-tie architecture, the core neurons are ALPNs, and the peripheral neurons are categorized as either feed-in or feed-out [50, 51]. Feed-in neurons, including olfactory sensory neurons and ALLNs, serve as sensory information sources that project into the core neurons. As the core neurons, ALPNs mediate a set of highly conserved processes for sensory information processing, supporting the network robustness [50]. Feed-out neurons, including neurons in the mushroom body and lateral horn, facilitate the flexible adaptation to diverse inputs and execution of appropriate outputs [50, 52]. Such bow-tie architecture suggests better robustness and adaptability of the olfactory pathway in *Drosophila*.

The connectivity matrix of the neural clusters also captures the synaptic preferences of each neural group (Fig.3E, Fig.4). On the one hand, a single neural cluster exhibits one specific dominant synaptic type to other neural clusters (Fig.3E). Considering that all neural vector representations are initialized with the all-one vectors, BHGNN-RT can take full advantage of the connectivity information for the neural embeddings. For example, clusters C1 and C2 of ALPNs are inferred for GABAergic neurons with the dominant GABAergic synapses, while other ALPN clusters are mainly for cholinergic neurons (Fig.3E). Similar cases can be observed in LHLNs, LHCENTs, and other neural classes. The detailed element strength of the connectivity matrix can be examined in Supplementary Fig.5. On the other hand, specific connectivity preferences between pairwise neural clusters can be observed across the connectivity matrix. Taking DANs as an example, their nine neural clusters only display sparse intra-class connections (as the yellow box in Fig.3E), whereas they possess many interactions with neurons in the mushroom body, including MBINs, KCs, and MBONs. However, the DAN cluster C1 shows extra reciprocal connections with other neural classes, including ALPNs, LHLNs, and LHCENTs, which cannot be observed from other DAN clusters. This suggests that DAN cluster C1 may serve as a global regulator along the olfactory pathway.

**Figure 4.**
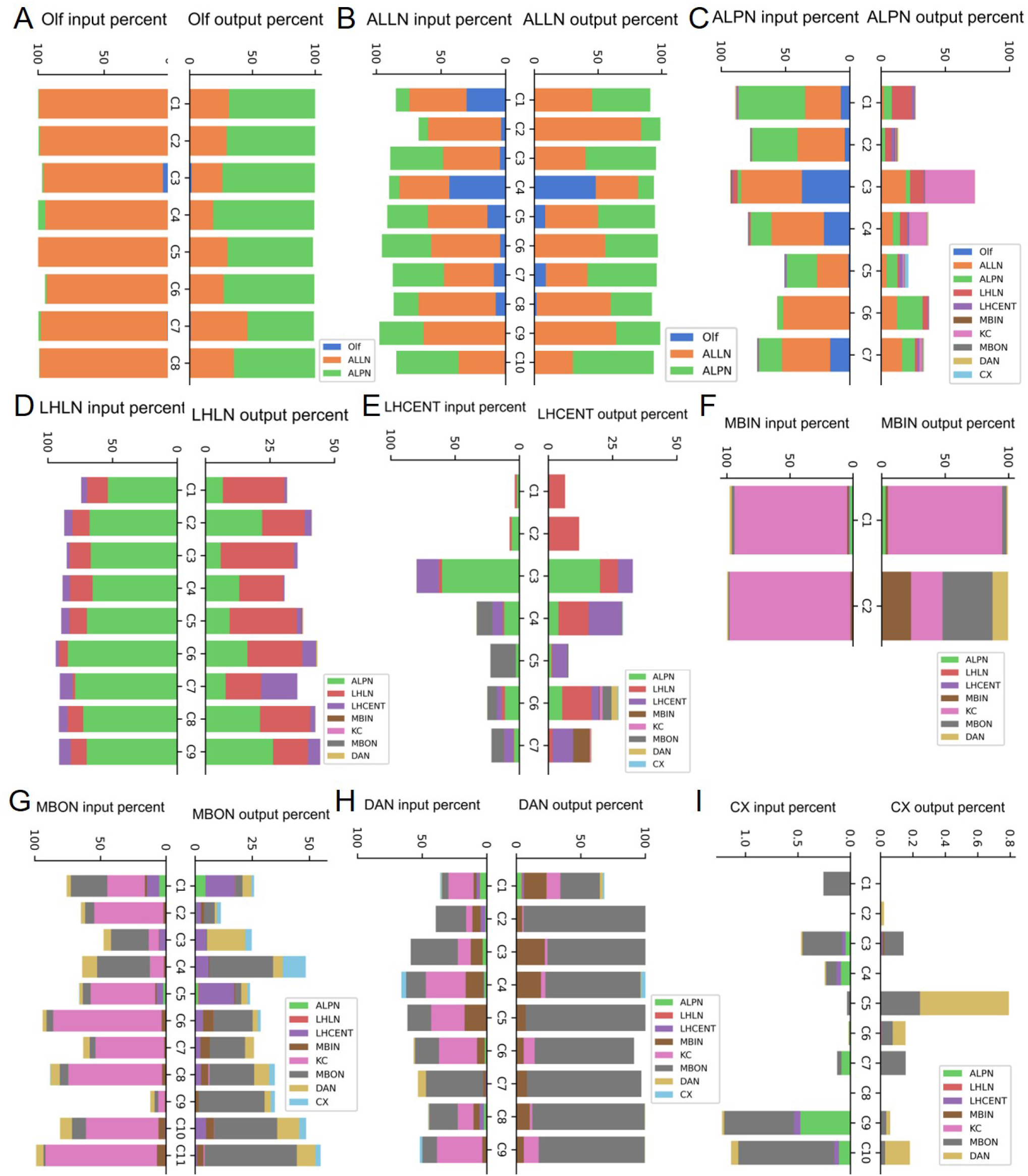
Connectivity profiles of the major neuronal classes along the olfactory pathway. The percentage of synaptic input and output of clusters of the major neural classes, when considering their connectivity with the major neuronal classes along the olfactory pathway. (A-I) denote the results for olfactory neurons (A), ALLNs (B), ALPNs (C), LHLNs (D), LHCENT (E), MBINs (F), MBONs (G), DANs (H), and central complex neurons (I).

### Bilateral symmetry of community architectures along olfactory pathway

The brain connectome in bilaterian organisms typically exhibits a lateralized architecture characterized by the presence of asymmetrical neural circuits predominantly localized in one hemisphere [53]. The olfactory system in *Drosophila* is also structured to efficiently process olfactory information through both ipsilateral and contralateral connections, which contributes to the robustness and precision of olfactory perception and behavior. In this section, we further investigate the bilateral connectivity patterns of neural clusters within the fly olfactory system.

The ipsilateral and contralateral connections along the olfactory pathway are depicted separately as Fig.5 and Supplementary Fig.7. Among them, these ipsilateral connections generally follow the connectivity patterns of the overall olfactory system (Fig.3A, Fig.5A). Olfactory receptor neurons in the antennal lobe send their axons (∼10^4^ synapses) to both the ipsilateral and contralateral ALLNs and ALPNs. The contralateral communication between bilateral antennal lobes is facilitated by the commissural interneurons, allowing for the integration of olfactory signals and enabling odor localization through the comparison of sensory information from two sides [54]. ALPNs further convert processed olfactory information to higher-order brain centers, specifically the lateral horn and MB calyx. Ipsilateral ALPNs predominantly project to the same-side LH and calyx (Fig.5), where they facilitate innate responses and associative learning, respectively. A subset of ALPNs also projects contralateral (∼10^2^ synapses to contralateral LHLNs, LHCETNS, and MBONs; ∼10^3^ synapses to contralateral KCs), enabling bilateral representation of odor information within these higher centers [11, 54]. In the MB, KCs receive predominant synaptic inputs from ipsilateral ALPNs (∼10^5^ synapses) and additional contralateral ALPN inputs (∼10^3^ synapses). The ipsilateral dominance of KC inputs supports lateralized processing, while contralateral connections enhance robustness and resilience to sensory perturbation. One thing worth noting is that MB only exhibits reciprocal connections (∼10^3^ synapses) within bilateral MBONs, while not in bilateral MBINs and KCs. Further, MBONs can receive contralateral projections from KCs (∼10^4^ synapses) and MBINs (∼10^3^ synapses). This balanced bilateral MBON connectivity ensures that olfactory cues, regardless of which hemisphere they are received from, can trigger consistent behavioral outcomes.

**Figure 5.**
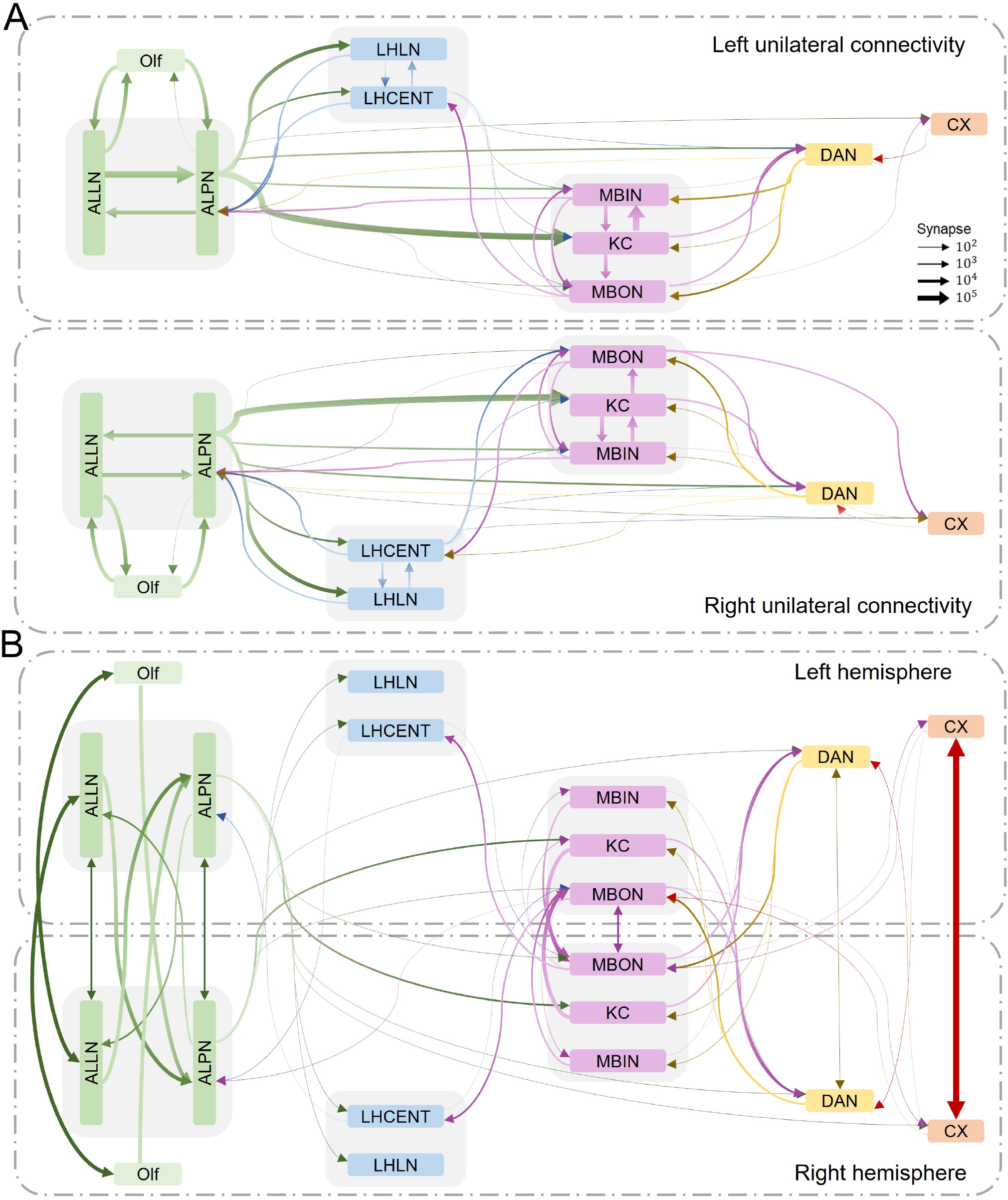
Analysis of symmetric connectivities along the olfactory pathway. (A) Unilateral connectivities among the major neuronal classes within unilateral hemispheres. (B) Contralateral connectivities among the major neural classes across two hemispheres.

We further investigate the connectivity profiles for neural clusters of this bilateral olfactory system. Considering the neuronal localization, the individual neural cluster in Fig.3E is separated into two subsets, each in one unilateral hemisphere. The corresponding connectivity matrix is visualized as Fig.6, which can be segmented based on ipsilateral connections (L→L, R→R) and contralateral connections (L→R, R→L) in the *Drosophila* adult brain. The matrix elements still represent the dominant synaptic type between pairwise neural clusters. Overall, the ipsilateral connectivity profiles in two hemispheres (L→L, R→R) are almost symmetric, which can also be observed in contralateral connections. Meanwhile, the contralateral connections are much sparser when compared with the ipsilateral connectivities. LHLNs, LHCENTs, MBINs, and KCs exhibit only ipsilateral neural connectivity in the *Drosophila* brain. This ipsilateral nature of these connections supports rapid, specialized neural processing and reduces the complexity of bilateral communication, enhancing the efficiency of response generation in olfactory-driven behaviors.

**Figure 6.**
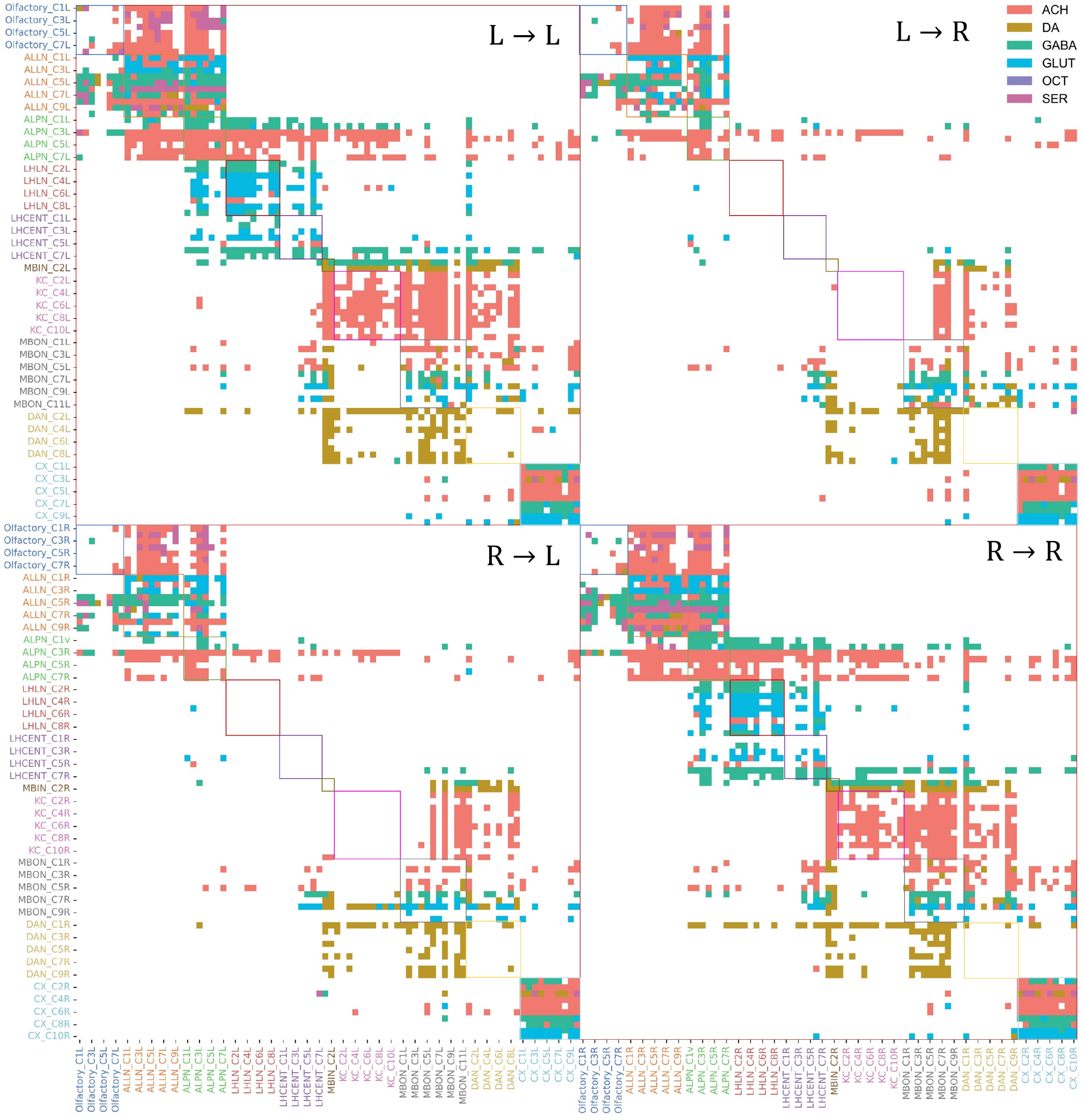
Network organization across multisensory pathways. (A) The distribution of shortest path lengths between neuron pairs in the largest strongly connected component (SCC) and weakly connected component (WCC) of FlyWire. Here, the SCC contains 119,770 neurons, 89.25% of all neurons, while the WCC covers 132,490 neurons, approximately 98. 73% of all neurons. (B) Number of neurons involved along multisensory pathways, with a path length ranging from 1 to 14. The starting points with PL=0 are the sensory neurons themselves. (C) Number of synapses involved in each layer along these six sensory pathways, based on the results of panel B. (D) Distributions of neurons with different neurite patterns along these sensory pathways, with the path length ranging from 0 to 5. To simplify panel D, we only show the distribution when the neural proportion in each neurite pattern is greater than 1%. The vertical axis corresponds to these neurite patterns.

Furthermore, the connectivity profiles of neural clusters can provide us with some hints about their specific functional roles. Taking the ALLN cluster C9 for example, it mainly receives contralateral connections from olfactory receptor neurons, while receiving little information from ipsilateral olfactory neurons. This might indicate that this ALLN neural group is specialized to enhance the comparison and integration of bilateral olfactory signals. Also, ALPN cluster C3 displays a prominent connectivity tendency to the contralateral LH and MB neural groups, via cholinergic projections. This may support more comprehensive odor coding and enhance the reliability of memory formation and behavioral output. Another obvious example is the DAN cluster C1, possessing ipsilateral connections to ALPNs, LHLNs, and LHCENTs that cannot be observed from other clusters. This unique ipsilateral wiring allows these specific DANs to rapidly modulate olfactory-driven behaviors based on sensory input without requiring cross-hemisphere processing, which may expedite responses to salient stimuli.

Concerning the bilateral symmetry of the fly olfactory system, we also investigate the difference in structural connectivities of contralateral neural groups. The percentages of synaptic connections for the major neural clusters are visualized in Fig.7 and Supplementary Fig.8. The majority of these neural clusters in two hemispheres, including olfactory sensory neurons (Fig.7A, D), ALLNs (Fig.7B, E), ALPNs (Fig.7C, F), MBINs (Fig.7G, J), KCs (Fig.7H, K), LHLNs (Supplementary Fig.7A-B), and LHCENTs (Supplementary Fig.7C-D), show symmetry connectivity profiles. Of course, slight differences in neural connectivity remain among these neural clusters. For example, Cluster C4 of olfactory receptor neurons in the right hemisphere owns about 10% input from the ipsilateral connections of ALPNs, which can not be observed in the left hemisphere. In Fig.7, the most prominent difference lies in the connectivity patterns of MBONs. The original MBON cluster C11 (Fig.3E) only includes one single neuron in the left hemisphere, hence we derived 11 MBON clusters in the left brain and 10 clusters in the right. The most obvious difference of MBON clusters happens among C7-C10 clusters, where neurons display diverse connectivity profiles. For example, MBON C10 in the left hemisphere receives almost 50% input from KC in the right hemisphere, while MBON C10 in the right obtains little information from KCs. This indicates the asymmetry within the bilateral MB regions also exists for the function of associative learning and memory.

**Figure 7.**
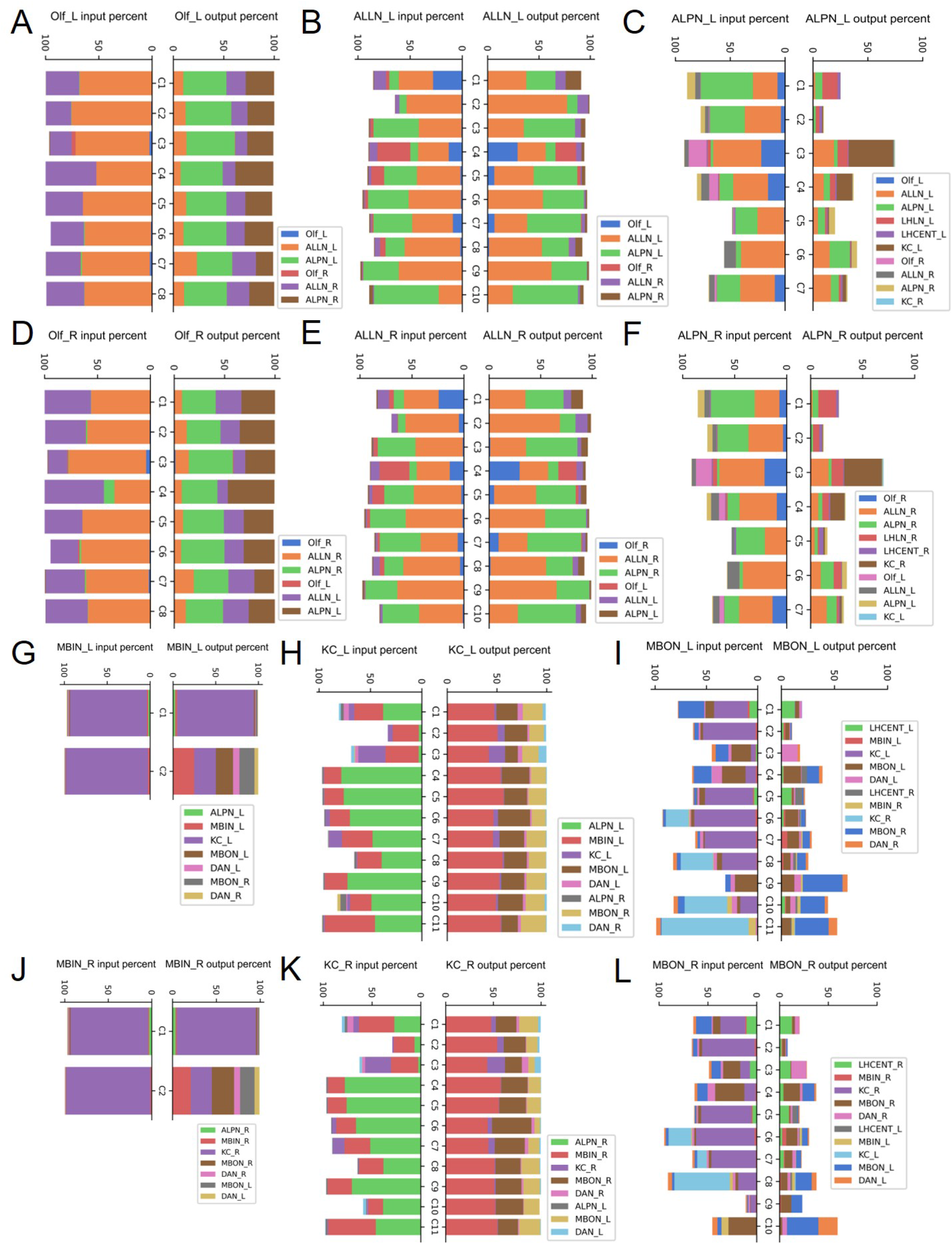
Connectivity component of neuronal clusters within both hemispheres. The percentage of synaptic input and output of clusters of the major neural classes, when considering the neural clusters within two hemispheres. (A-C, G-I) denote the results of the olfactory neurons, ALLNs, ALPNs, MBINs, KCs, and MBONs in the left brain. (D-F, J-L) represent the results of these neural classes in the right brain. Other neural classes are shown in the supplementary figures.

In this section, our comprehensive connectivity analysis of the olfactory pathway elucidates the complex bilateral integration and neurotransmitter-specific connections that facilitate robust sensory processing and adaptive behavioral responses, providing deeper insights into the organizational principles of the Drosophila olfactory system.

### Hierarchical organization across multisensory pathways

In the *Drosophila* brain, sensory information is hierarchically processed to produce effective behavioral outcomes. From the network perspective, we can observe a structured architecture that enables both quick adaptation to environmental cues and high-level decision-making processes. In this section, our analysis provides insights into how different layers of sensory integration contribute to both rapid neural responses and complex decision-making.

Based on the *Drosophila* brain connectome (FlyWire dataset v783), we first examined its largest connected component in which each neuron can be linked to another one via one or more paths. The largest weakly connected component (WCC) comprises 132,490 neurons, representing 98.73% of all neurons that are interconnected when directionality is ignored. In contrast, 119,770 neurons, or 89.25% form the largest strongly connected component (SCC) via directed pathways. The distribution of shortest path length between pairwise neurons within these two components, as shown in Fig.8, indicates that most neuron pairs are connected within four or five steps, suggesting close interconnectivity throughout the network. Besides, the maximum shortest distance, or the network diameter, is 14, which reflects the extensive reachability of neurons across the brain.

**Figure 8.**
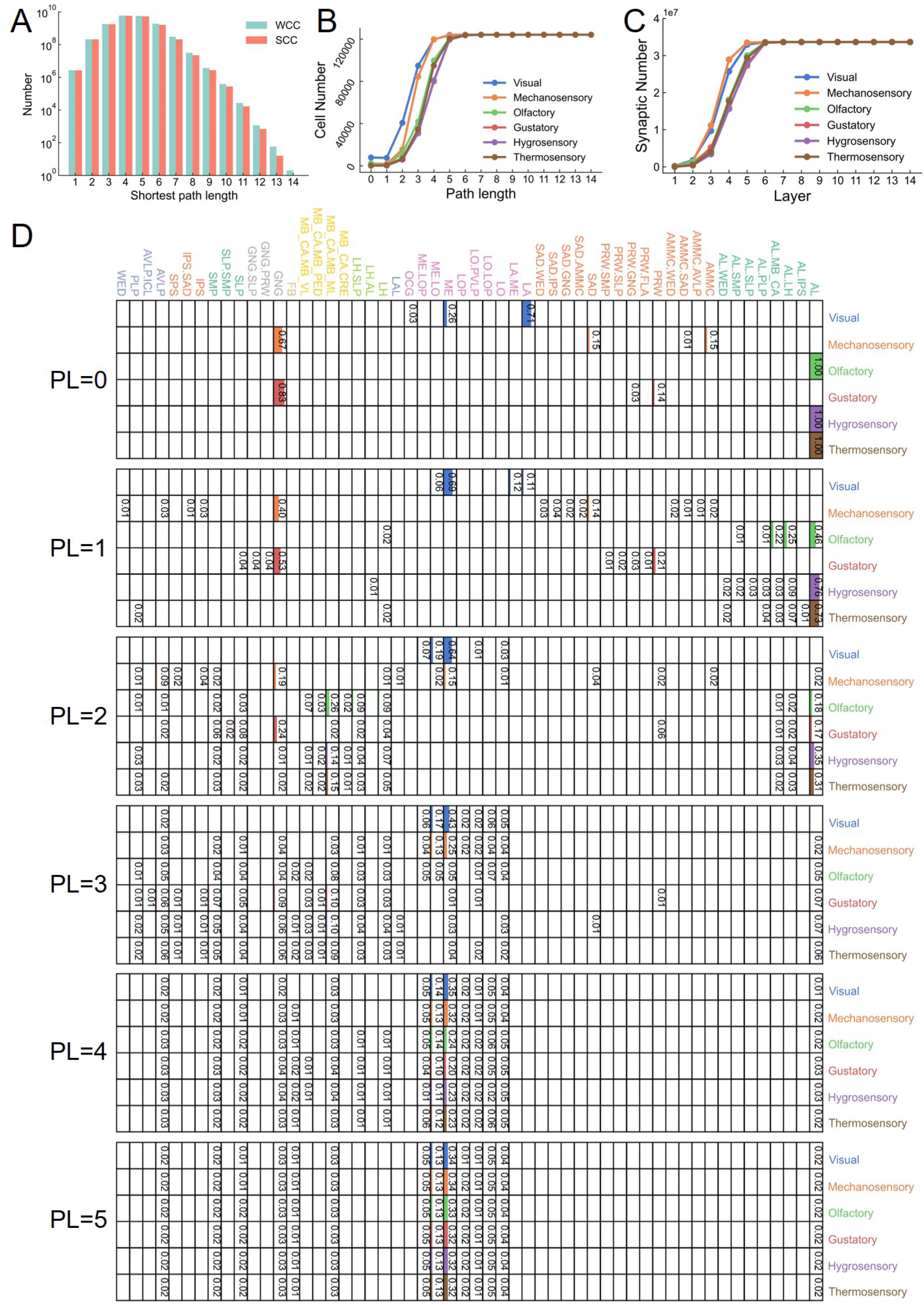
Network organization across multisensory pathways. (A) The distribution of shortest path lengths between neuron pairs in the largest strongly connected component (SCC) and weakly connected component (WCC) of FlyWire. Here, the SCC contains 119,770 neurons, 89.25% of all neurons, while the WCC covers 132,490 neurons, about 98.73% of all neurons. (B) Number of neurons involved along these six sensory pathways, with the path length ranging from 1 to 14. The starting points with PL=0 are the sensory neurons themselves. (C) Number of synapses involved in each layer along these six sensory pathways, based on the results of panel B. (D) Neuron distributions of different neurite patterns for these sensory pathways with the path length ranging from 0 to 5. To simplify panel D, it only displays the distribution when neural proportion with individual neurite pattern is larger than 1%. The vertical axis lists these neurite patterns. AL, AL.IPS, AL.LH, AL.MB_CA, AL.PLP, AL.SLP, AL.SMP, AL.WED are mainly located in Antennal Lobe. AMMC, AMMC.AVLP, AMMC.SAD, AMMC.WED, PRW, PRW.FLA, PRW.GNG, PRW.SLP, PRW.SMP, SAD, SAD.AMMC, SAD.GNG, SAD.IPS, SAD.WED are mainly in Periesophageal Neuropils. LA, LA.ME, LO, LO.LOP, LO.PVLP, LOP, ME, ME.LO, ME.LOP, OCG are in Optic Lobe. LAL is in lateral complex. LH, LH.AL, LH.SLP are mainly located in the Lateral Horn. MB_CA.CRE, MB_CA.MB_ML, MB_CA.MB_PED, MB_CA.MB_VL are in Mushroom Body. FB is in the central complex. GNG, GNG.PRW, GNG.SLP are in the Gnathal ganglia. SLP, SLP.SMP, SMP are in Superior Neuropils. IPS, IPS.SAD, SPS are in Ventromedial region. AVLP, AVLP.ICL, PLP, WED are in Ventrolateral Neuropils.

Along the typical sensory pathways, we examined the involvement of neurons and synapses with path lengths ranging from 1 to 14 (Fig.8B-C). Starting from sensory neurons at a path length (PL) of 0, we observed that the number of neurons and synapses gradually increases, particularly within the range of 1 to 5 steps. After five steps, the number of involved neurons and synapses begins to converge across different pathways, indicating a consistent pattern of sensory integration at higher levels. This convergence suggests that sensory inputs are efficiently integrated within six hops to almost all neurons in the fly brain, consistent with previous findings on the hierarchical nature of sensory processing for coherent perception and behavior [1, 5].

We further analyzed the distribution of neurons with different neurite patterns across sensory pathways for path lengths between 0 and 5 (Fig.8D). The neurite pattern of a single neuron is categorized by the major location of its dendrites and axon. These neurite patterns can provide insights into the key regions involved in processing specific sensory inputs and highlight anatomical distinctions across different sensory pathways. In Fig.8D, only neurons with the proportion of a specific neurite pattern exceeding 1% are displayed. The olfactory, hygrosensory, and thermosensory sensory neurons predominantly exist in the antennal lobe, with the major dendrites and axons all in the antennal lobe. At PL=1, a larger proportion of downstream neurons of hygrosensory and thermosensory neurons are still located in the AL (76% and 73%, respectively), compared to the direct downstream neurons (46%) of olfactory sensory neurons. This may suggest that weaker sensory signals, such as humidity and temperature, require further amplification and preprocessing at early stages. Conversely, olfactory neurons show greater involvement of mushroom body neurons (26%, with neurite pattern of MB_CA.MB_ML) at an early stage (PL = 2), indicating that olfactory information can be delivered faster to MB for information processing and associative learning than other sensory modalities. As the sensory signals propagate through the network, other sensory pathways increasingly interact with neurons in the optic lobe, suggesting cross-modal integration with visual information. With additional steps (PL=4 and 5), the neuron distributions along these six sensory pathways become more similar, indicating convergent processing at higher levels. Similarly, mechanosensory and gustatory also demonstrate a consistent pattern, originating from the gnathal ganglia (GNG) and converging with other brain regions as they advance through the network.

In sum, our analyses illustrate that distinct sensory neurons are linked with specific neurite patterns, reflecting specialized processing centers for each sensory modality. Strong similarities are evident among olfactory, hygrosensory, and thermosensory pathways, particularly at early processing stages. As sensory signals travel through the network, cross-modal interactions increase, such as the consistent neurite patterns in the mushroom body and higher brain regions. This hierarchical organization enables multisensory integration, where information from distinct sensory streams merges to create a new, unified response that is different from the individual sensory inputs. Our findings emphasize that efficient connectivity, specialized processing centers, and layered sensory integration contribute to sophisticated multisensory processing in the *Drosophila* brain.

### Interaction between multisensory neurons and the olfactory pathway

Based on the above analysis, we further explore how other sensory modalities interact with the olfactory system to support complex behaviors. In *Drosophila* adult, the typical sensory neurons consist of visual, mechanosensory, olfactory, gustatory, hygrosensory, and thermosensory neurons [11, 45, 55]. These sensory neurons respond to diverse environmental cues, including optical, auditory, odor, taste, humidity, and temperature information. Extensive multisensory integration patterns are observed throughout multiple interconnected pathways with distinct path lengths, such as olfactory, thermosensory, and hygrosensory pathways [5, 21]. The multisensory interactions are mainly facilitated through neural projections that converge on the lateral horn and mushroom body, which serve as key centers for associative learning and innate behavior [42]. To make it simple, here we mainly discuss the direct interactions between multisensory neurons and the primary olfactory pathway.

Among these multisensory neurons, the visual neuron group, with almost 8000 cells, occupies the majority of the sensory population, while only very few hygrosensory and thermosensory neurons exist (Fig.9A). Relative to the number of these sensory neurons, the neural centralities, including degree centrality and eigenvector centrality, show opposite distributions (Fig.9B). Hygrosensory and thermosensory neurons display higher centrality compared with other neural types. Takemura found that hygrosensory and thermosensory neurons establish connections not only within their sensory domains but also with some olfactory circuits, suggesting that they play an essential role in the integration of environmental conditions with behavioral responses [42]. The high centrality metrics just reflect their importance in modulating key adaptive behaviors, such as regulating humidity or temperature preferences, which are crucial for maintaining homeostasis and survival.

**Figure 9.**
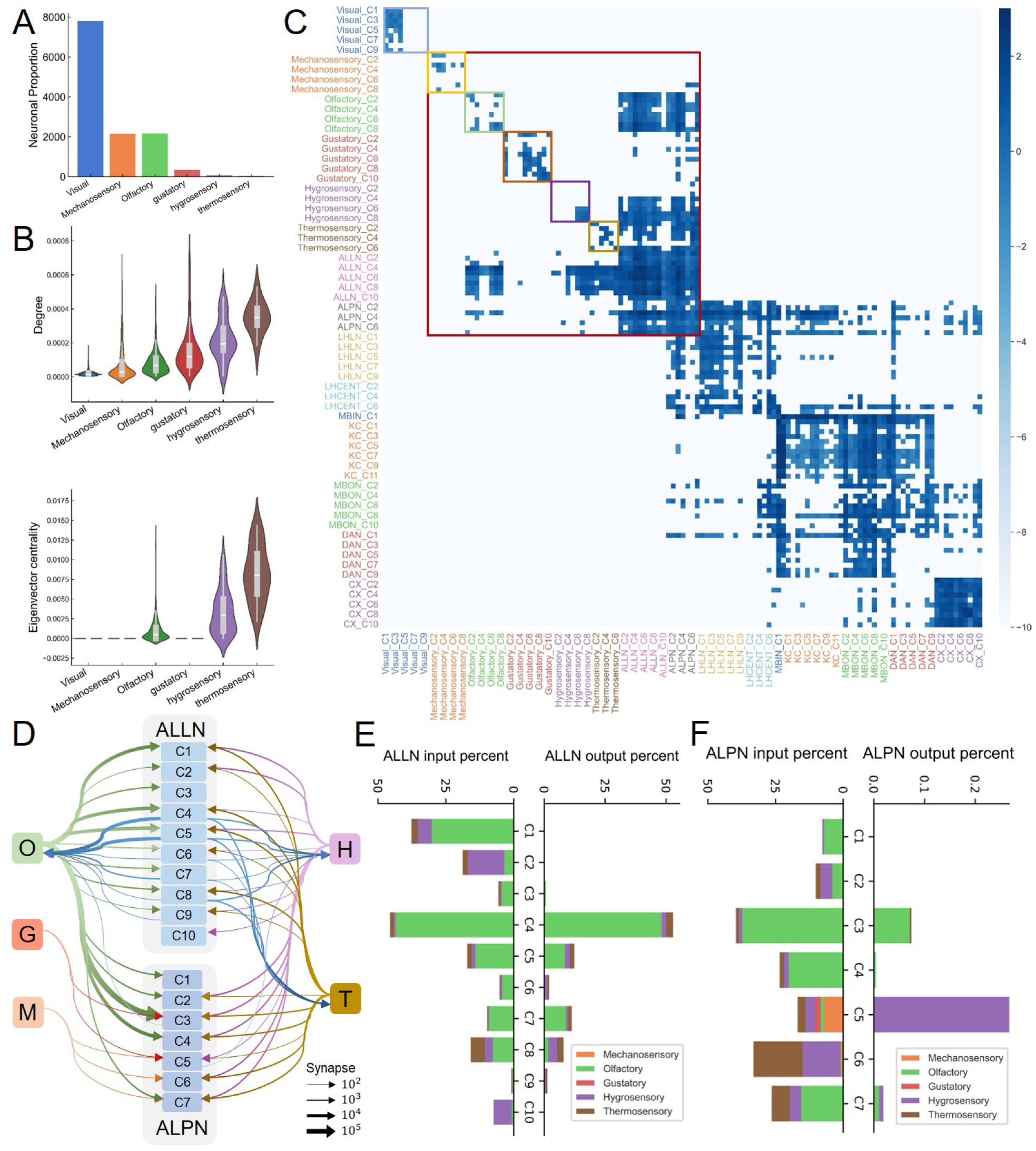
Interactions between the olfactory pathway and other sensory neurons. (A) Neural numbers of the main neuronal classes for six main sensory regions, including visual, mechanosensory, olfactory, gustatory, hygrosensory, and thermosensory neurons. (B) Centrality organizations of these main neuronal classes. (C) The connectivity matrix of clusters for these six sensory neuronal groups. The cluster results are inferred based on neural vector representation via BHGNN-RT, following similar procedures in Fig.3B-D. The element in the logarithmic connectivity matrix is the synaptic strength between pairwise clusters, normalized by the neuronal numbers of these two corresponding clusters. (D) A schematic of the synaptic connectivity among sensory neurons and ALLN clusters and ALPN clusters. (E-F) The percentage of synaptic input and output of ALLN and ALPN clusters from different sensory modalities.

Based on the embedding results, we generated the clustering results for these sensory neurons (Fig.9C, Supplementary Fig.9) and examined the connectivity profiles of the corresponding neural clusters. In Fig.9C, the connectivity matrix displays the logarithmic connection strengths between pairwise neuron clusters. Individual squares mark the intraclass connectivities of these sensory neurons. Meanwhile, the predominant synaptic type of these pairwise neural clusters is depicted in Supplementary Fig.10A. The intraclass connectivities of sensory neurons are much sparser than those of other neurons, such as ALLNs, ALPNs, and others. Meanwhile, these six types of sensory neurons do not possess direct projections between each other. However, except for visual neurons, all other sensory neurons have direct connections to neurons in the AL region (Fig.9C). Mechanosensory neurons exhibit weak connections to ALPNs, which start from the Mechanosensory cluster C7. Gustatory neurons also primarily project to ALPNs, while remaining slight connections to one ALLN cluster (ALLN C4). As for hygrosensory and thermosensory neurons, they all have strong connections to the majority of ALLNs and ALPNs, similar to the connectivity patterns of olfactory receptor neurons. Meanwhile, olfactory, hygrosensory, and thermosensory neurons also receive stronger feedback information from ALLNs, especially ALLN clusters C4-C9.

The observed differences among AL neuron clusters in their connectivity patterns reflect the distinct functional roles in sensory processing and integration. We depict a more intuitive schematic of synaptic connectivity among the clusters of sensory neurons, ALLNs, and ALPNs and the percentage of the relevant synaptic connections (Fig.9D-F). ALLN clusters C1-C2, which receive unidirectional input from olfactory, hygrosensory, and thermosensory neurons, are likely involved in specialized processing of environmental cues for rapid, context-specific responses (Fig.9E). In contrast, ALLN clusters C3 and C10, which obtain specific sensory modalities, appear to specialize in processing distinct types of sensory input with minimal integration, thus preserving the fidelity of each sensory signal. ALLN clusters C4-C5 and C7-C8, showing strong reciprocal connections with olfactory, hygrosensory, and thermosensory neurons, serve as key hubs for integrating diverse sensory inputs. This highlights their importance in cross-modal integration and fine-tuning of sensory representations within the antennal lobe. Similar connectivity patterns can also be observed in the ALPN clusters. ALPN clusters display connectivity preferences for integrating different combinations of sensory modalities. For example, ALPN cluster C3, which primarily receives direct input from olfactory receptor neurons, specializes in encoding olfactory signals with high fidelity. ALPN cluster C6, receiving inputs mainly from hygrosensory and thermosensory neurons, is crucial for integrating environmental humidity and temperature, allowing *Drosophila* to adapt to specific microclimatic conditions. Cluster C5, which handles almost all sensory modalities, likely plays a broader role in generating a multisensory representation of the environment, essential for context-dependent decision-making. Besides, the fact that all ALPN clusters generate few feedback projections suggests a primary feedforward architecture optimized for efficient signal transmission and minimal signal distortion during sensory input processing.

Overall, the interaction between multisensory neurons and the olfactory pathway emphasizes the sophisticated way in which *Drosophila* integrates different types of sensory information to produce adaptive behavior. The direct convergence of olfactory, mechanosensory, gustatory, hygrosensory, and thermosensory neurons in the antennal lobe ensures that a single sensory cue is processed within a broader sensory context, learning to produce innate adaptive responses.

## Discussion

This study provides a comprehensive analysis of the structural connectivity and hierarchical organization of multisensory pathways in the *Drosophila* brain. Our analysis of the FlyWire dataset reveals significant details about the network properties of the *Drosophila* adult brain connectome. The degree distributions indicate varied connectivity patterns among different neurons. Some of them act as hubs with extensive connections, while others have more specialized roles with limited synaptic partners. The observed neuronal and synaptic variabilities highlight the complexity of biological neural circuits, which is critical for maintaining network stability and efficient information processing.

To effectively leverage the unique properties of neuronal profiles for message-passing and to capture network heterogeneity, we developed a network embedding method called BHGNN-RT. Extensive experiments demonstrated its efficacy in the task of unsupervised clustering for directed, heterogeneous networks. BHGNN-RT achieved superior performance compared to other baseline methods, with the highest silhouette scores by grouping similarly labeled nodes more accurately. Further analysis revealed that BHGNN-RT effectively identified distinct spatial and functional groups among Kenyon cells, illustrating their roles in sensory integration and signal transmission within the mushroom body.

Our examination of the olfactory pathway uncovered the community architecture among various neural classes, including neurons in the antennal lobe, lateral horn, and mushroom body. Fig.3 visualizes these connectivity patterns, highlighting the significant roles of GABAergic and cholinergic neurons in sensory modulation and signal relay within the antennal lobe. Meanwhile, ALPNs and MBONs possess direct cholinergic projections targeting neurons in the central complex, thereby shortening the communication pathways from primary sensory regions to higher brain centers. Moreover, specific dopaminergic neuron clusters exhibit distinct connectivity preferences to neurons in the lateral horn and mushroom body, underscoring their functional roles in learning and memory. This structured community architecture facilitates stepwise integration and refinement of sensory inputs, enhancing the complexity and accuracy of information processing. Additionally, our analysis of bilateral connectivity patterns in the olfactory pathway (Fig.5-Fig.7) highlights significant intra-hemisphere connections within neural classes such as LHLNs, LHCENTs, MBINs, and KCs. In contrast, sparse contralateral connections in other regions support bilateral coordination, ensuring integrated olfactory processing across hemispheres. Besides, although neuronal connections along bilateral olfactory pathways show overall symmetry, specific asymmetries are also present, especially in some neural clusters of MBONs.

The interactions between multisensory neurons and the olfactory pathway demonstrate extensive integration across sensory modalities in the *Drosophila* brain. The connectivity matrix in Fig.9 illustrates how olfactory, gustatory, mechanosensory, hygrosensory, and thermosensory neurons directly converge in the antennal lobe, where diverse connectivity preferences shape information processing. The multisensory integration within the antennal lobe is essential for forming cohesive sensory experiences at early stages and guiding appropriate behaviors. Furthermore, we investigated the hierarchical organization of multisensory pathways throughout the *Drosophila* brain connectome. The hierarchical structure supports efficient sensory processing, facilitating rapid signal transmission and comprehensive integration. Sensory inputs can be efficiently integrated within six hops to almost all neurons in the fly brain. According to the analysis of neurite patterns, these major sensory pathways exhibit strong similarities of neuronal organizations at higher layers. This further supports the concept of functional segregation and integration within the brain.

This work represents the first application of connectivity-based clustering to a biological neural network through graph representation learning. It provides a detailed map of hierarchical community architecture in the *Drosophila* brain, offering valuable insights into neural circuit organization and the integration of multisensory information. Our findings highlight the importance of balanced connectivity preferences in maintaining network stability and efficient signal processing. Further research should explore the dynamic aspects of the brain connectome, examining how connectivity and neural activity change in response to environmental stimuli and learning tasks. Extending our analysis to other model organisms may yield comparative insights into the fundamental principles governing neural organization and multisensory integration.

## Materials and methods

### Datasets

#### FlyWire database for Drosophila adult brain connectome

According to the latest version of the FlyWire dataset (v783), which maps the entire *Drosophila* brain, 139,255 neurons have been proofread, and 34,153,566 synapses labeled within the standard brain template (FAFB) [45]. Given the current incompleteness of the connectome and its annotations, we focused on using neural data for which there is greater confidence, including information regarding neuron type, class, cell type, neurite pattern, and cell body location. These attributes are relatively complete and provide a reliable basis for our analysis.

In addition, we merged synapses between neuron pairs that shared the same neurotransmitter type, eliminating possible duplication in pairwise neural connections. This step helped refine our dataset by reducing redundancy and highlighting unique functional connections. Our inspection of FlyWire v783 revealed that only 134,191 neurons have annotated connections, with a total of 2,831,350 unique connections mapped. Despite the incomplete nature of the dataset, focusing on well-annotated neural features allows us to form meaningful insights into the structure and connectivity of the brain.

#### Benchmark datasets for evaluating BHGNN-RT

We evaluated the performance of the proposed network embedding method, BHGNN-RT, using several benchmark datasets and compared it with other state-of-the-art algorithms. The six public datasets used in our evaluation include Cora, Cora_ml, CiteSeer, CiteSeer_full, Amazon_CS, and Amazon_photo [56, 57, 58]. These datasets represent directed heterogeneous networks, encoding directed relationships between nodes that represent different entities. The Cora, Cora_ml, CiteSeer, and CiteSeer_full datasets consist of nodes representing research articles, with edges denoting citation relationships between them. Each article is described by a bag-of-words feature vector that captures the presence or absence of specific terms from the dictionary. Amazon_CS and Amazon_photo represent the Amazon co-purchase network, where nodes correspond to products and edges indicate co-purchase relationships between different goods. The feature vectors for these datasets are derived from bag-of-words extracted from product reviews, providing insights into the associations between products based on customer preferences. In these datasets, node types correspond to the ground-truth classes, allowing for effective validation of the embedding quality.

### Fitting for distribution functions

To extract the distribution pattern for the degree distribution (Fig.1A-B) and the synaptic weight distribution (Fig.1C-D), we applied the following five common probability distributions to select the best fitting distribution (Table 1). The lower fitting bound of the degree distribution is denoted as *x*_min_ and *C* is the normalization constant satisfying 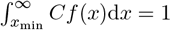 in Table 1 [59]. For the degree distributions and synaptic weight distributions, we set *x*_min_ = 1. The Akaike Information Criterion (AIC) is used to determine the best fit among all distribution candidates. Detailed fitting results are presented in Table 2–3 for the degree distributions and synaptic weight distributions.

**Table 1:** Probability density functions on [*x*_min_, ∞) used for fitting distribution patterns.

| Name | $C$ | $f(x)$ |
| --- | --- | --- |
| Power-law (Pareto) | $(\alpha - 1)x_{\min}^{\alpha-1}$ | $x^{-\alpha}$ |
| Gamma | $\frac{\lambda^{1-\alpha}}{\Gamma(1-\alpha, \lambda x_{\min})}$ | $x^{-\alpha} e^{-\lambda x}$ |
| Exponential | $\lambda e^{\lambda x_{\min}}$ | $e^{-\lambda x}$ |
| Weibull | $\beta \lambda e^{\lambda x_{\min}^{\beta}}$ | $x^{\beta-1} e^{-\lambda x^{\beta}}$ |
| log-normal | $\sqrt{\frac{2}{\pi \sigma^2}} [\operatorname{erfc}(\frac{\ln x_{\min} - \mu}{\sqrt{2}\sigma})]^{-1}$ | $\frac{1}{x} \exp[-\frac{(\ln x - \mu)^2}{2\sigma^2}]$ |

**Table 2:**
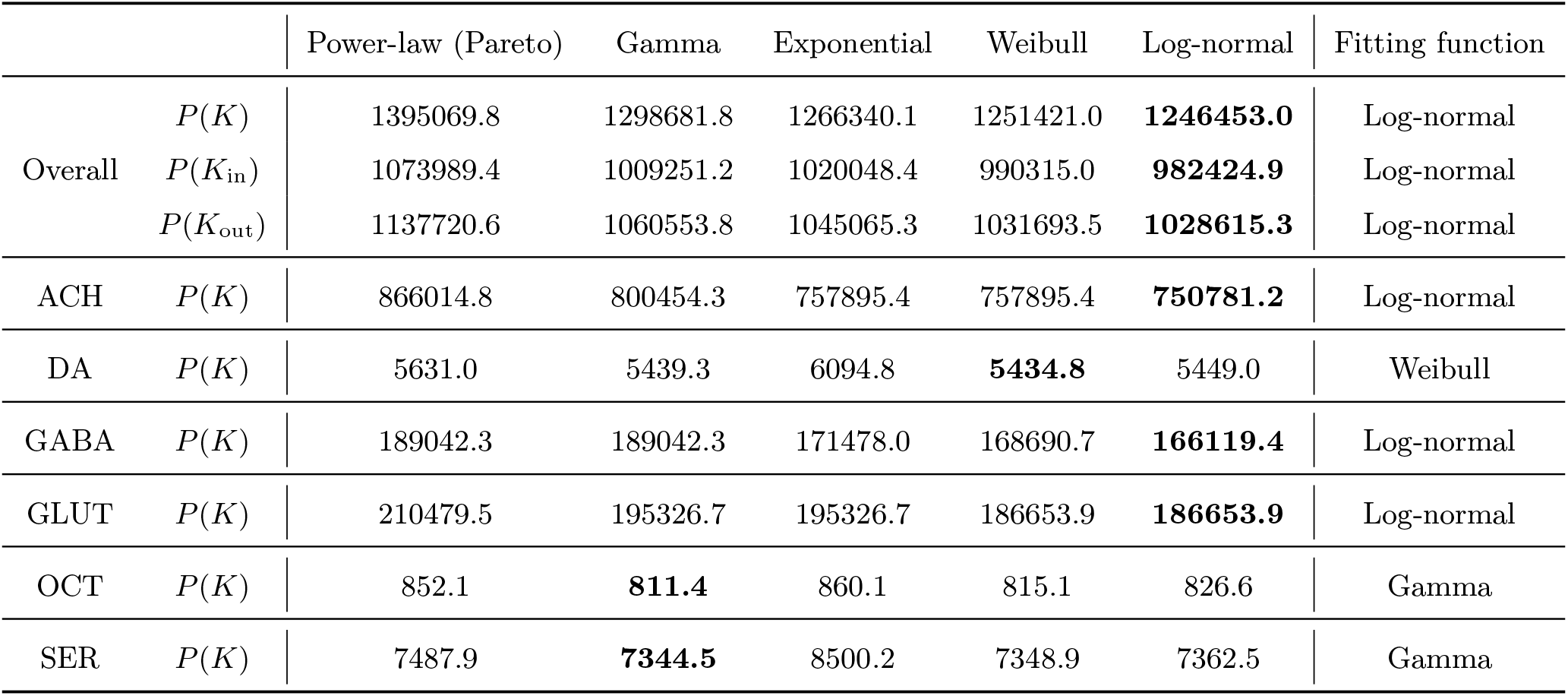
Model selection for degree distribution based on AIC.

|  |  | Power-law (Pareto) | Gamma | Exponential | Weibull | Log-normal | Fitting function |
| --- | --- | --- | --- | --- | --- | --- | --- |
| Overall | $P(K)$ | 1395069.8 | 1298681.8 | 1266340.1 | 1251421.0 | **1246453.0** | Log-normal |
| | $P(K_{\text{in}})$ | 1073989.4 | 1009251.2 | 1020048.4 | 990315.0 | **982424.9** | Log-normal |
| | $P(K_{\text{out}})$ | 1137720.6 | 1060553.8 | 1045065.3 | 1031693.5 | **1028615.3** | Log-normal |
| ACH | $P(K)$ | 866014.8 | 800454.3 | 757895.4 | 757895.4 | **750781.2** | Log-normal |
| DA | $P(K)$ | 5631.0 | 5439.3 | 6094.8 | **5434.8** | 5449.0 | Weibull |
| GABA | $P(K)$ | 189042.3 | 189042.3 | 171478.0 | 168690.7 | **166119.4** | Log-normal |
| GLUT | $P(K)$ | 210479.5 | 195326.7 | 195326.7 | 186653.9 | **186653.9** | Log-normal |
| OCT | $P(K)$ | 852.1 | **811.4** | 860.1 | 815.1 | 826.6 | Gamma |
| SER | $P(K)$ | 7487.9 | **7344.5** | 8500.2 | 7348.9 | 7362.5 | Gamma |

### Statistical analysis of network properties in the *Drosophila* brain

The *Drosophila* adult brain connectome of *Drosophila* represents a comprehensive map of neural circuitry, encompassing the intricate web of neural connections and their corresponding synaptic weights. Investigating its statistical characteristics is a crucial step in unraveling the structural principles and complexities of the *Drosophila* neural network. In this subsection, we shall mainly examine three main network properties: degree distribution, synaptic weight distribution, and network heterogeneity.

#### Degree distributions

The degree distribution of a network, which describes the number of connections that each node has, is essential for understanding its structural characteristics [60]. In a directed graph, the distributions in-degree and out-degree can reveal important insights into how information flows within the network [21].

Here, the degree distributions are depicted via complementary empirical cumulative distribution functions (CECDF), plotted on logarithmic axes. CECDF is defined as 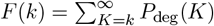, which sums the probability for a node with a degree *K* larger than *k*. This function decreases monotonically along *k* with the limits where *F* (0) = 1 and *F* (*k*_*max*_ + 1) = 0. Meanwhile, plots on logarithmic axes are more helpful for visualizing their heavy-tailed distributions.

We investigated the overall degree distribution and canonical neural degree distributions (shown as Supplementary Fig.1A-B). To further uncover the network topology, these distribution patterns were examined by fitting with commonly-used distribution functions, including power-law (Pareto), gamma, exponential, Weibull, and log-normal distributions. Detailed information is described in the Table 1. Furthermore, Akaike’s information criterion (AIC), a widely used method for model selection that balances model fit and complexity, is applied to determine the best fit among all distribution candidates.

The AIC formula is defined as AIC = 2*k* − 2*LL*, where *k* is the number of estimated parameters and *LL* is the log-likelihood for the model [61]. The detailed fitting results are illustrated in Table 2, where a smaller value represents a better fitting.

According to the fitting results, we found that the log-normal distribution provides the best fit for the overall degree, in-degree, and out-degree distributions (Supplementary Fig.1A). This indicates that while most neurons have a moderate degree, a significant number of neurons have a very high degree and a few neurons have a shallow degree. Considering the modular organization in *Drosophila* brain connectome, neurons within modules are highly interconnected, while connections between modules are less dense but still follow a hierarchical pattern. This log-normal distribution can emerge from these hierarchical processes where high-level hub neurons have exponentially more connections than low-level ones. It follows the mechanism of preferential attachment in network analysis, where the rich become richer. In addition, the in-degree and out-degree distributions usually display completely different patterns in real-world networks [62, 63]. However, similar patterns of neural in-degree and out-degree distributions occur in *Drosophila* brain connectome (Supplementary Fig.1A), which reflects the intricate balance of neural connections.

Meanwhile, we investigated the degree distributions of the canonical neuronal types, including cholinergic, GABAergic, glutamatergic, dopaminergic, octopaminergic, and serotonergic neurons (Supplementary Fig.1B, Table 2). The major neuron types (cholinergic, GABAergic, and glutamatergic) all fit log-normal distribution very well Supplementary Fig.1B left), while the minority obey Weibull or gamma distributions (Supplementary Fig.1B right). The maximal degree of GABAergic neurons is an order of magnitude higher than others. The variation in degree distribution among different neuronal types highlights the diversity of their connectivity patterns, which can directly affect the functional roles of these neurons. Cholinergic, GABAergic, and glutamatergic neurons are the principal excitatory or inhibitory neurons. Their broad and variable connectivity, aligned with log-normal function, may be required for rapid synaptic transmission, inhibitory control, and excitatory signaling. While for dopaminergic, octopaminergic, and serotonergic neurons, Weibull or gamma distribution may reflect their specialized, modulatory roles with more selective connectivity patterns.

#### Synaptic weight distributions

Synaptic weight, which reflects differences in the strength and importance of connections [10], directly affects the efficacy and dynamics of neural communication. Investigating the synaptic weight distribution enables us to determine whether the network exhibits a few strong connections amidst many weak ones or just a more uniform weight distribution [64]. Such analysis is vital for modeling neural processes, as it affects how signals propagate through the network and how the brain integrates and processes information.

Supplementary Fig.1C shows the distribution of synaptic weights *P* (*w*) with its best fit of log-normal distribution (Table 3). When comparing the exponential fit, a heavy tail can be observed, which indicates that these strong synaptic connections cannot be overlooked. Supplementary Fig.1D presents the synaptic weight distributions for the canonical neuronal types. Similar to the result of Supplementary Fig.1B, the synaptic weight distributions of the major neuron types, including cholinergic, GABAergic, and glutamatergic neurons, all follow the log-normal distribution, while others fit gamma or Weibull distributions (Table 3). Fitting with log-normal distribution is consistent with the results of excitatory cells in rat cortex [65], which presents the log-normal fit instead of others. Meanwhile, the peak occurs when *w* = 5 among all these distributions. It implies the majority of neurons work in harmony with few synaptic connections. These differences in synaptic weight distributions reflect variations in how different neuronal types modulate synaptic strength.

**Table 3:**
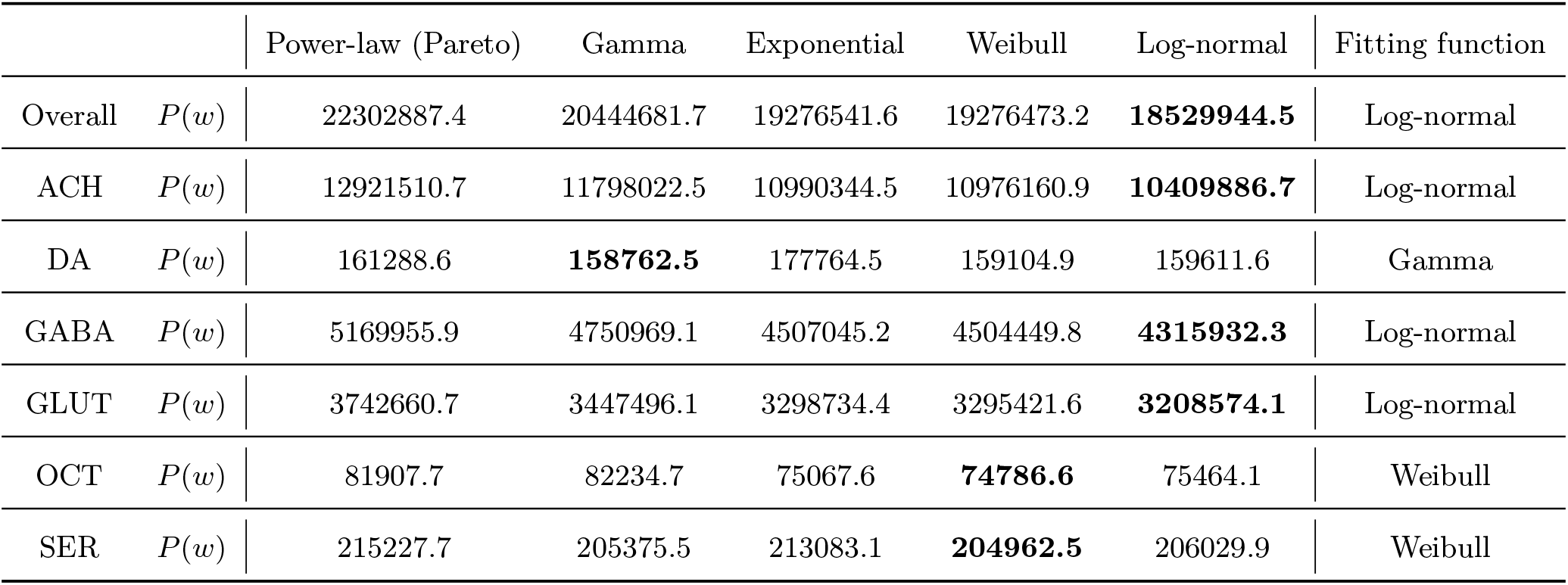
Model selection for distribution of synaptic strength based on AIC.

#### Network heterogeneity: neuron and synapse diversity

Network heterogeneity refers to the diversity of the types of neurons and synapses [29]. In the context of the *Drosophila* brain connectome, the heterogeneity of its neural classes and synaptic types is depicted in Supplementary Fig.1E-F. Supplementary Fig.1E shows the proportion of various neuron types, with cholinergic neurons (over 60%) being the most prevalent. Cholinergic, GABAergic, and glutamatergic neurons cover at least 87.9% of all neurons in *Drosophila* brain Supplementary Fig.1E. By the way, the cell type of the remaining 10.9% neurons, which are labeled as unclear in Supplementary Fig.1F, has not yet been clarified. We further explored the synaptic diversity, based on the neurotransmitter type (Supplementary Fig.1F left) and pre-/post-synaptic neuronal types (Supplementary Fig.1F right). The left panel follows the consistent pattern in Supplementary Fig.1E, and the right panel of Supplementary Fig.1F demonstrates that the significant dominance of cholinergic, GABAergic, and glutamatergic connections remains. Investigations on network heterogeneity highlight the intricate organization within the *Drosophila* brain connectome.

The traditional doctrine, “one neuron, one neurotransmitter”, held that synaptic communication between two neurons occurs via releasing a single chemical transmitter [66]. However, recent research suggests that neurons can release more than one specific neurotransmitter during neural communications [66, 67]. According to the neurotransmitter identifications in FlyWire, we illustrate the distribution of neurotransmitters released from individual neural types as Supplementary Fig.1G-H. Considering the complexity of neurotransmitter co-release, we mainly consider one predominant neurotransmitter in a single connection. Here, a connection is defined as a group of synapses with the same neurotransmitter between pairwise neurons. Among Supplementary Fig.1G-H, each type of neuron may release six distinct neurotransmitters. This phenomenon makes any neurons remarkably versatile, which can exert excitatory, inhibitory, and modulatory influences under different circumstances. Of course, the primary neurotransmitter released from one type of neuron is still consistent with the corresponding neural type (Supplementary Fig.1G), with the highest average synaptic strength (Supplementary Fig.1H).

Analysis of network properties collectively underscores the complexity and specialization within this directed, weighted, and heterogeneous graph, *Drosophila* brain connectome. This complexity would further hinder our understanding of hierarchical structure and functions in the biological neural circuits. In the following sections, our embedding method generates vector representations for all neurons in the FlyWire and attempts to infer their connectivity-based community structures within the *Drosophila* brain.

### Comparative analysis of embedding methods

We compared the performance of BHGNN-RT with other SOTA methods in the above-mentioned public datasets. These methods include DGI [40], DAEGC [68], GIC [69], JBGNN [27], and the variant of R-GCN (R-GCN-v) [28]. The comparative analysis in our previous work [37] demonstrates significant performance achieved by BHGNN-RT across all datasets. BHGNN-RT exceeds its performance over other baselines, with a higher clustering accuracy, ranging from 4.5% to 19.3%. It is promising that our proposed method allows each node stronger access to the structural properties of global connectivity. For a more intuitive comparison, we leveraged the t-distributed stochastic neighborhood embedding (t-SNE) method to reveal the relevant embedding patterns on the Cora dataset [37]. Corresponding embedding results were evaluated using SIL scores, a metric to quantify the quality of the clusters generated. BHGNN-RT achieves the highest SIL score of 0.477, much higher than other baseline scores, which indicates that node representations produced by BHGNN-RT can better separate same-labeled nodes into the same group. Therefore, our method improves unsupervised clustering quality when capturing more comprehensive nodal connectivity profiles and graph-level structural properties.

### Neuronal centrality in the connectome

To evaluate the topological importance of neurons, we mainly considered four centrality measures in this work, including degree centrality, eigenvector centrality, hub centrality, and authority centrality. Degree centrality is simply equivalent to the node degree in a network, where *C*_*D*_(*v*_*i*_) equals *k*_*i*_ for a node *v*_*i*_. Eigenvector centrality is a measure that accounts for the quality and quantity of the nodal connections. It considers the degrees of both a node *v*_*i*_ and its neighbors. Eigenvector centrality of the node *v*_*i*_ is measured as *C*_*E*_(*v*_*i*_) = *x*_*i*_, where *x*_*i*_ is the *i*-th element of the eigenvector *x* corresponding to the largest eigenvalue *λ*_1_ of the adjacency matrix ***A***. A node with a high value of *C*_*E*_ means it is connected to many other nodes or many other nodes with high eigenvectors. Similar to the eigenvector centrality, the hub and authority of a node corresponding to the *i*-th element of the dominant eigenvectors of the matrices ***AA***^*T*^ and ***A***^*T*^ ***A***, respectively. Normally, a node with a high authority centrality means that it is pointed to by many nodes with high hub centralities, and a node with a high hub centrality means that it points to many nodes with high authority centralities.

## Acknowledgments

We thank Dr. Hokto Kazama for his kind discussion of our analytical work and Dr. Hiroshi Kohsaka for his valuable comments on the paper revision. This work was supported by JSPS KAKENHI Grant Number JP22H00510.

## Conflict of interest

The authors declare no competing interests.

## Supplementary Figure

**Supplementary Figure 1:**
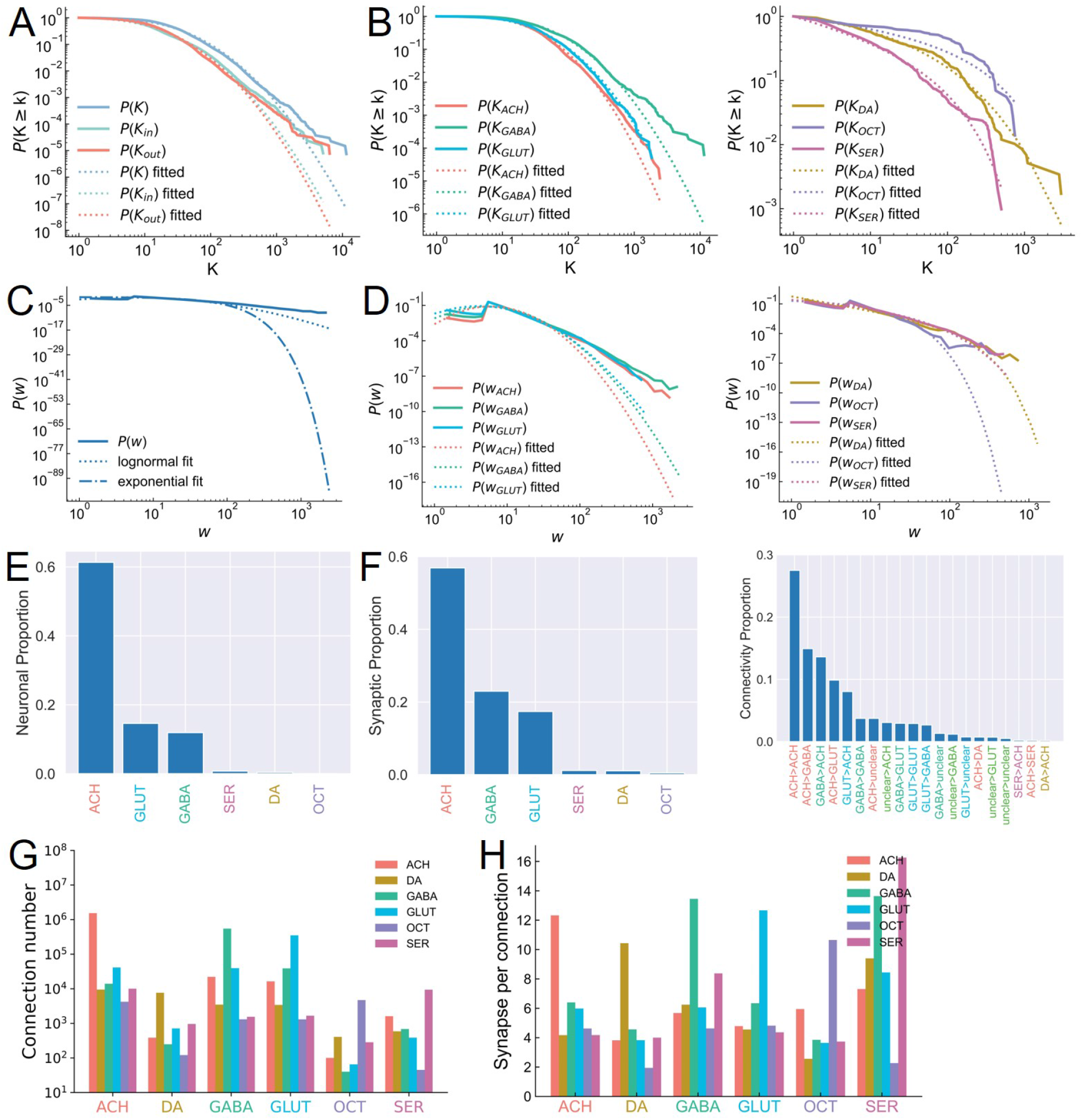
Network properties of the directed, weighted and heterogeneous graph (FlyWire dataset). (A) Degree, in-degree, and out-degree distributions of all neurons in FlyWire and their best fits. (B) Degree distributions of six main canonical neurons, including cholinergic (ACH), GABAergic (GABA), Glutamatergic (GLUT), Dopaminergic (DA), Octopaminergic (OCT), and serotonergic (SER) neurons, and their best fits. (C-D) Synaptic weight distribution for all neurons and their types. (E-F) Network heterogeneity is displayed when considering neuron diversity (E) and synaptic diversity (F). The left panel in F is based on neurotransmitters, while the right panel is based on pre-/post-synaptic neuronal type. (G-H) Network heterogeneity is quantified when considering the diversity of neurotransmitters released from one specific neural type. (G) Distributions of connection numbers with different neurotransmitters (ACH, DA, GABA, GLUT, OCT, SER) from one specific neural type. (H) The average synaptic strength among the distribution of connection number in (G).

**Supplementary Figure 2:**
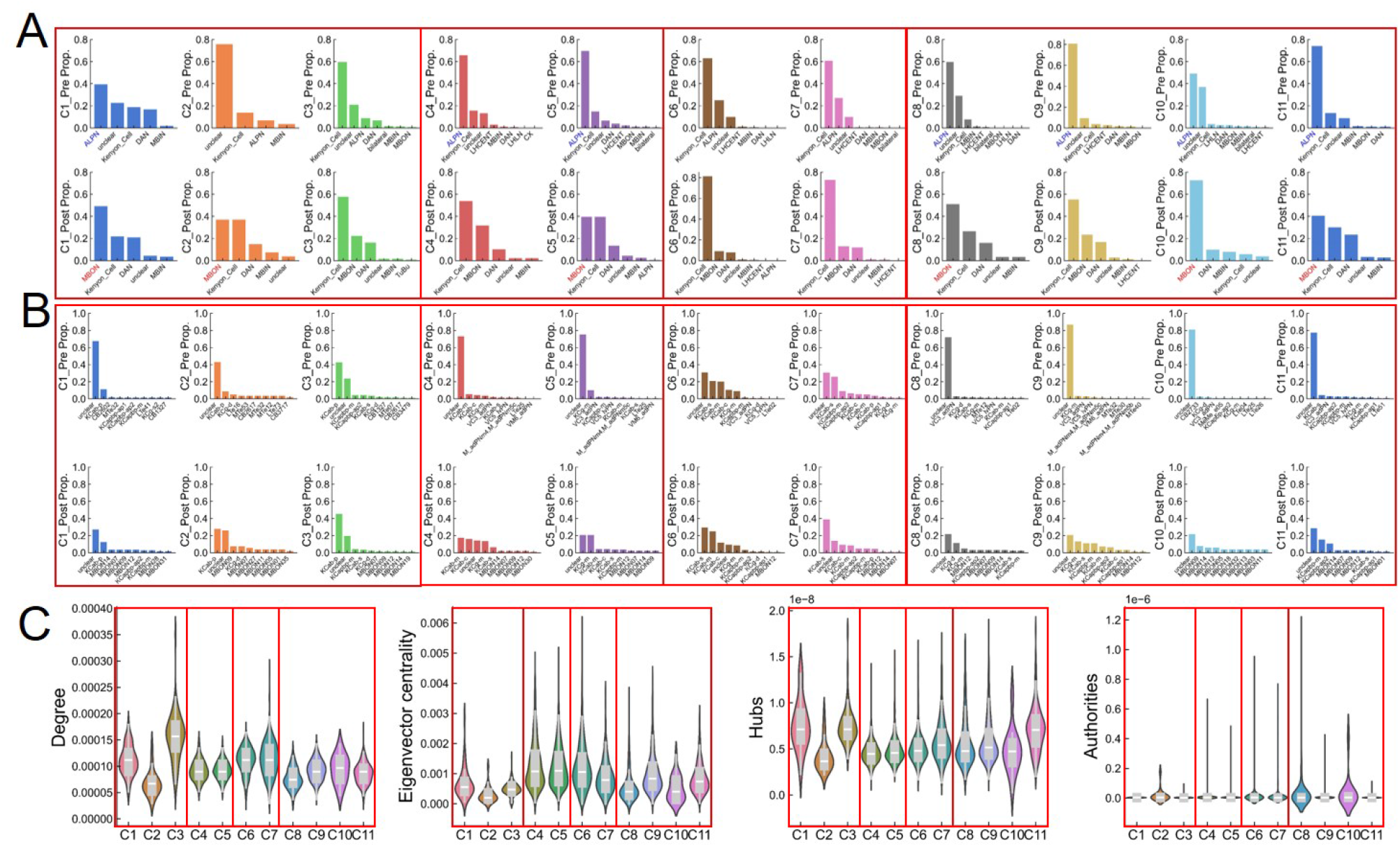
Connectivity patterns and quantification of KC clusters. (A) Distribution of presynaptic/postsynaptic neural types for individual clusters. (B) Distribution of presynaptic/postsynaptic neural classes for individual clusters. (C) Distribution of importance for each cluster, quantified by four main centralities, including 1) degree centrality; 2) eigenvector centrality; 3) hub centrality; and 4) authority centrality.

**Supplementary Figure 3:**
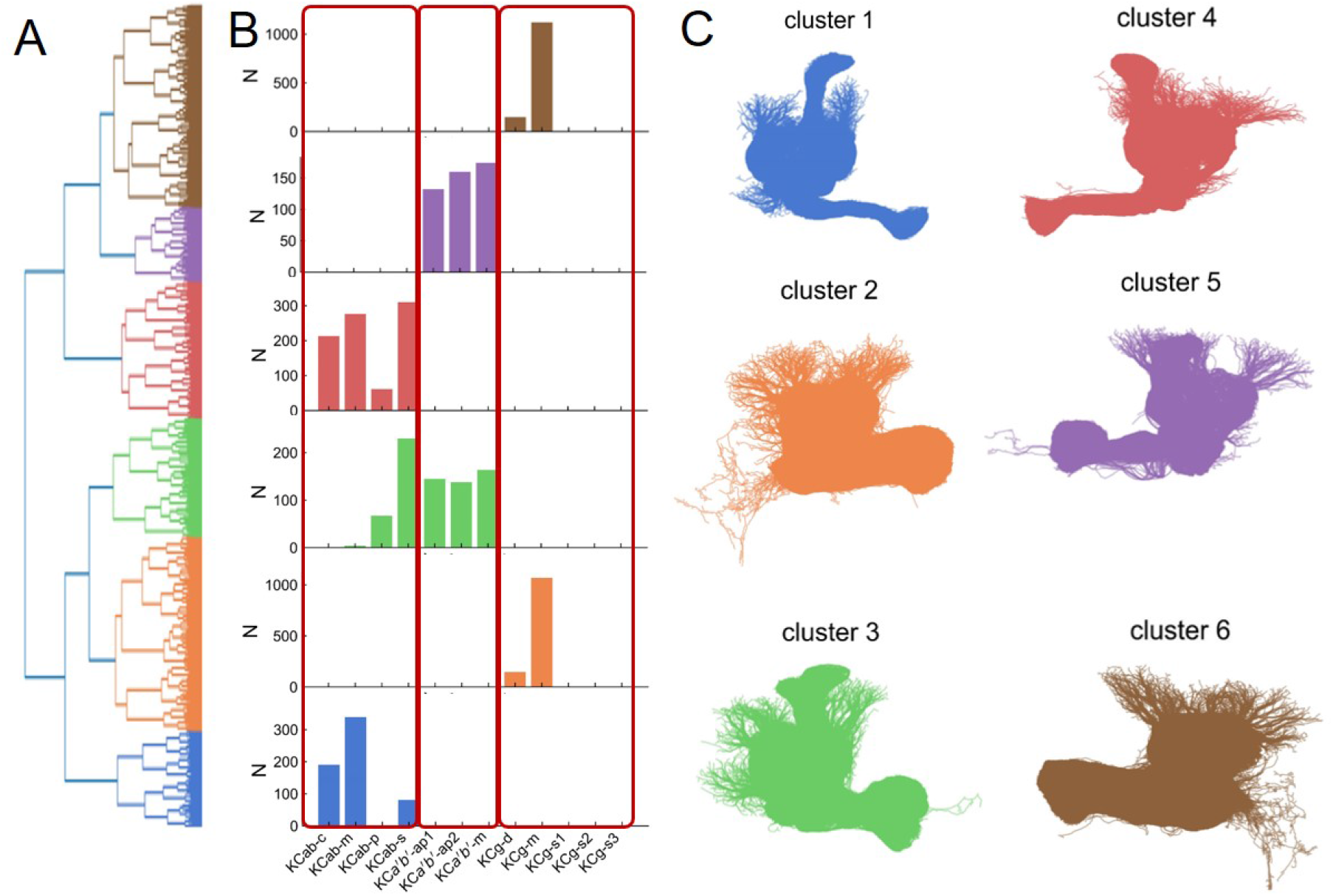
Morphological clustering for Kenyon Cells based on NBLAST. (A) Hierarchical clustering of KCs based on their morphological similarity calculated via NBLAST. (B) Distributions of cell type for these six neuronal groups based on the ground truth. KC *α, β* lobes include four main types (KCab-c, KCab-m, KCab-p, KCab-s), KC *α*^*′*^, *β*^*′*^ lobes include three main types (KCapbp-ap1, KCapbp-ap2, KCapbp-m), and KC *γ*^*′*^ lobe includes five main types (KCg-d, KCg-m, KCg-s1, KCg-s2, KCg-s3). (C) Spatial organizations of individual neuronal groups within KCs.

**Supplementary Figure 4:**
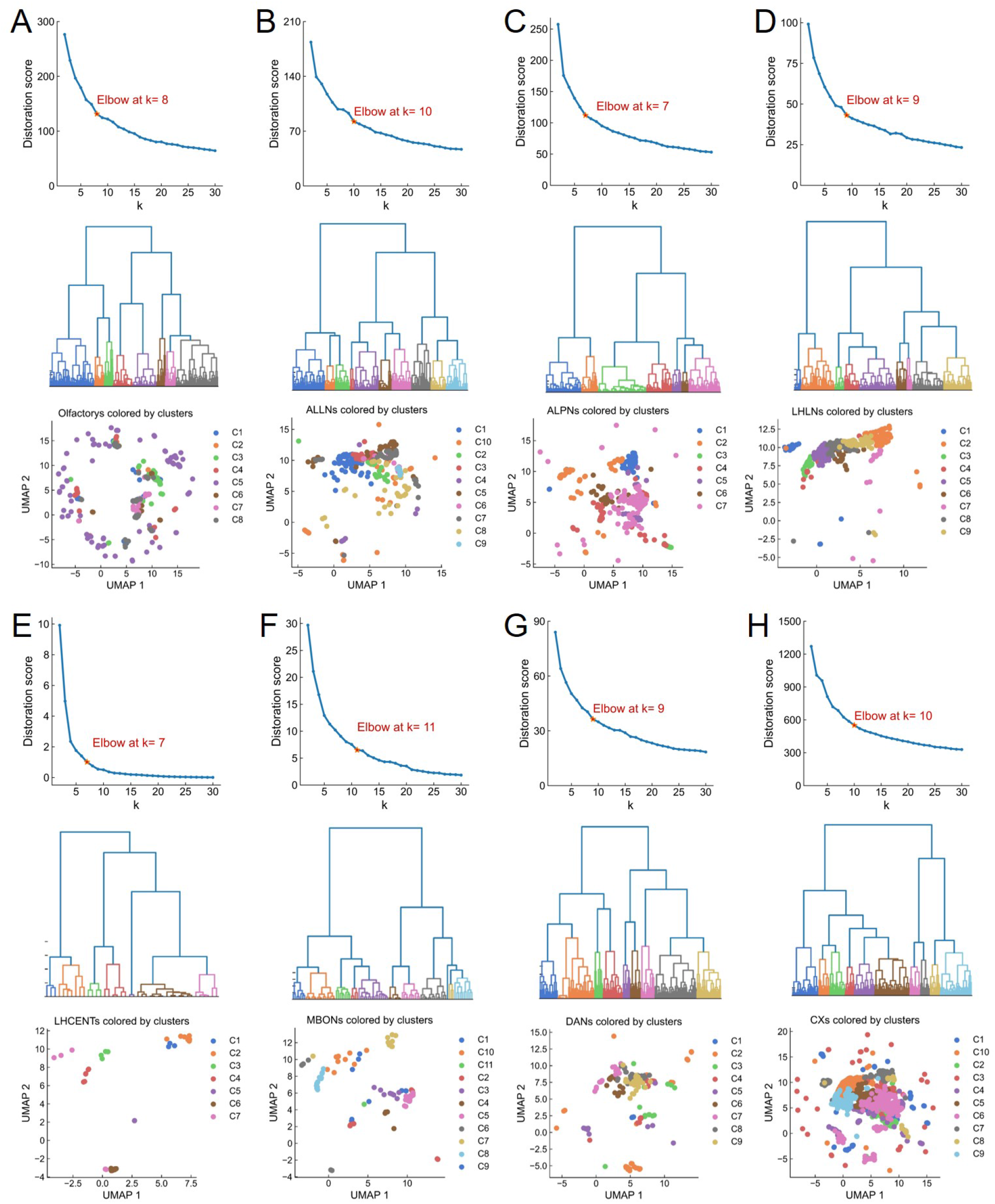
Hierarchical clustering results for the main neuron classes along the olfactory pathway. The elbow method is applied to determine the ideal number of neuronal clusters (first column), based on vector representations via BHGNN-RT. Hierarchical clustering of neuronal classes marked by colors of corresponding numbers (Second column). The UMAP visualization of vector representations for each neuronal class (Third column). These neuronal classes include olfactory sensory neurons (A), antennal lobe local neurons (ALLNs) (B), antennal lobe projection neurons (ALPNs) (C), lateral horn local neurons (LHLNs) (D), mushroom body output neurons (MBONs) (E), dopaminergic neurons (DANs) (F), and central complex neurons (CXs). Here, we did not present the clustering processing for mushroom body input neurons (MBINs), because there are only four giant MBINs in the *Drosophila* adult brain. 40

**Supplementary Figure 5:**
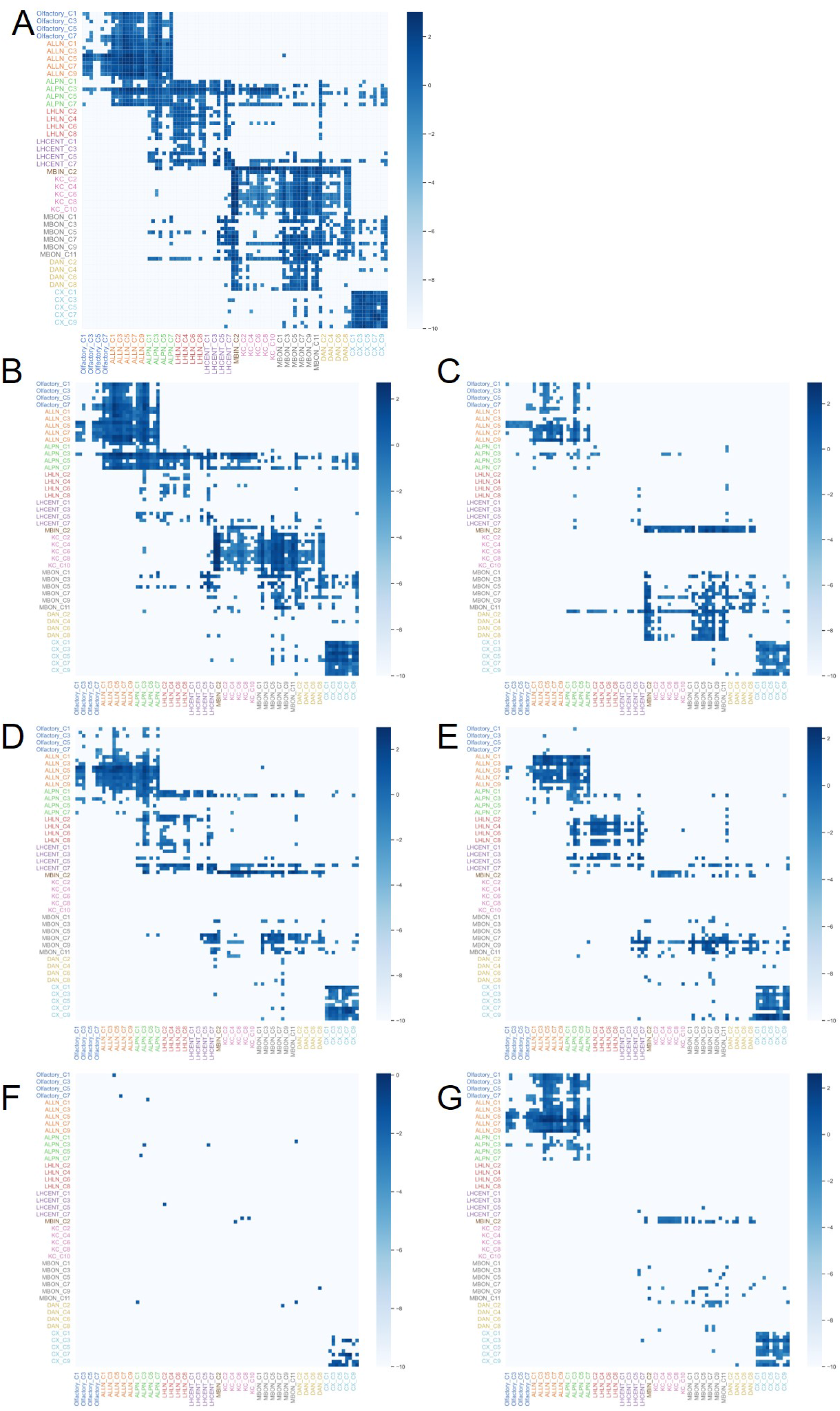
Connectivity matrix for neuronal clusters along the olfactory pathway. (A) The connectivity matrix of clusters for these eight nine neuronal classes. The cluster results are inferred based on neural vector representation via BHGNN-RT, following similar procedures in Fig.2B-C. The element in the connectivity matrix is the logarithmic synaptic strength between pairwise clusters, which is normalized by the neuronal numbers of these two corresponding clusters. The right panel is for the dominant connection type with a connection strength threshold of 5, according to the comparison across panels B-G. (B-G) The connectivity matrix of clusters is solely for synapses with unique neurotransmitters, including ACH (B), DA (C), GABA (D), GLUT (E), OCT (F), and SER (G).

**Supplementary Figure 6:**
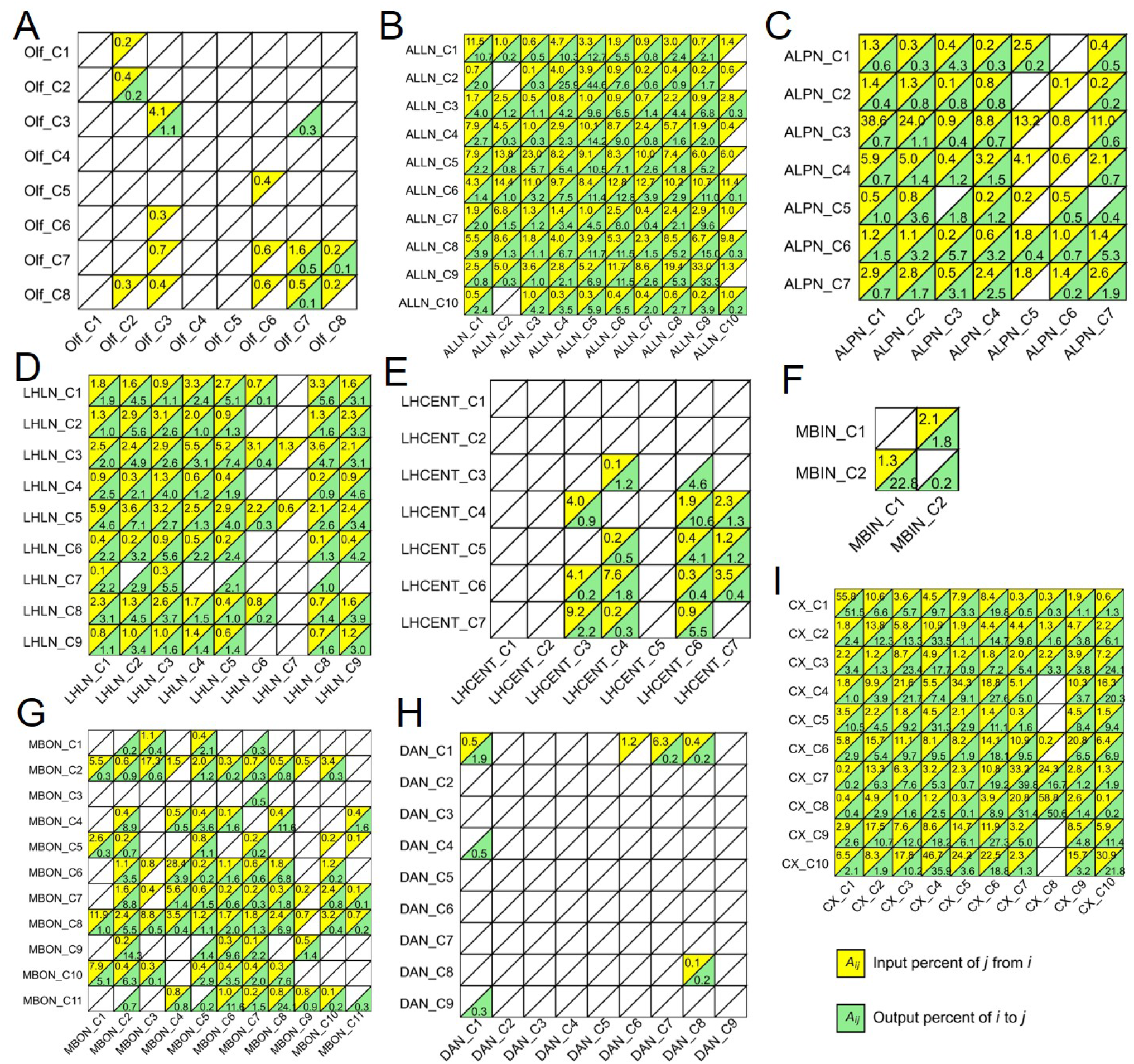
The percentage of synaptic input and output of inter-clusters for each neural class. They include olfactory neurons (A), ALLNs (B), ALPNs (C), LHLNs (D), LHCENTs (E), MBINs (F), MBONs (G), DANs (H) and CXs (I).

**Supplementary Figure 7:**
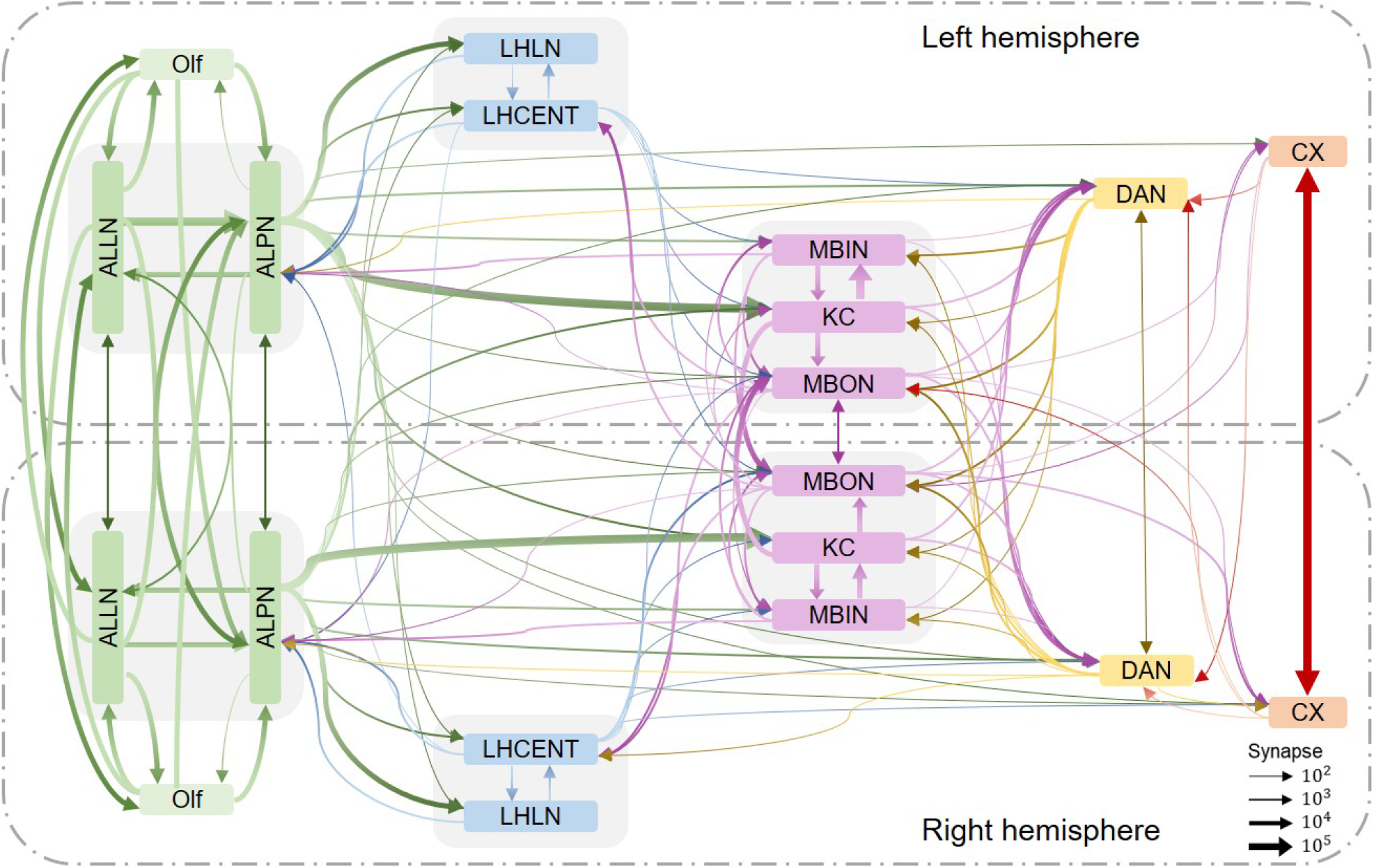
The complete connectivity of the olfactory system for ipsilateral and contralateral connections.

**Supplementary Figure 8:**
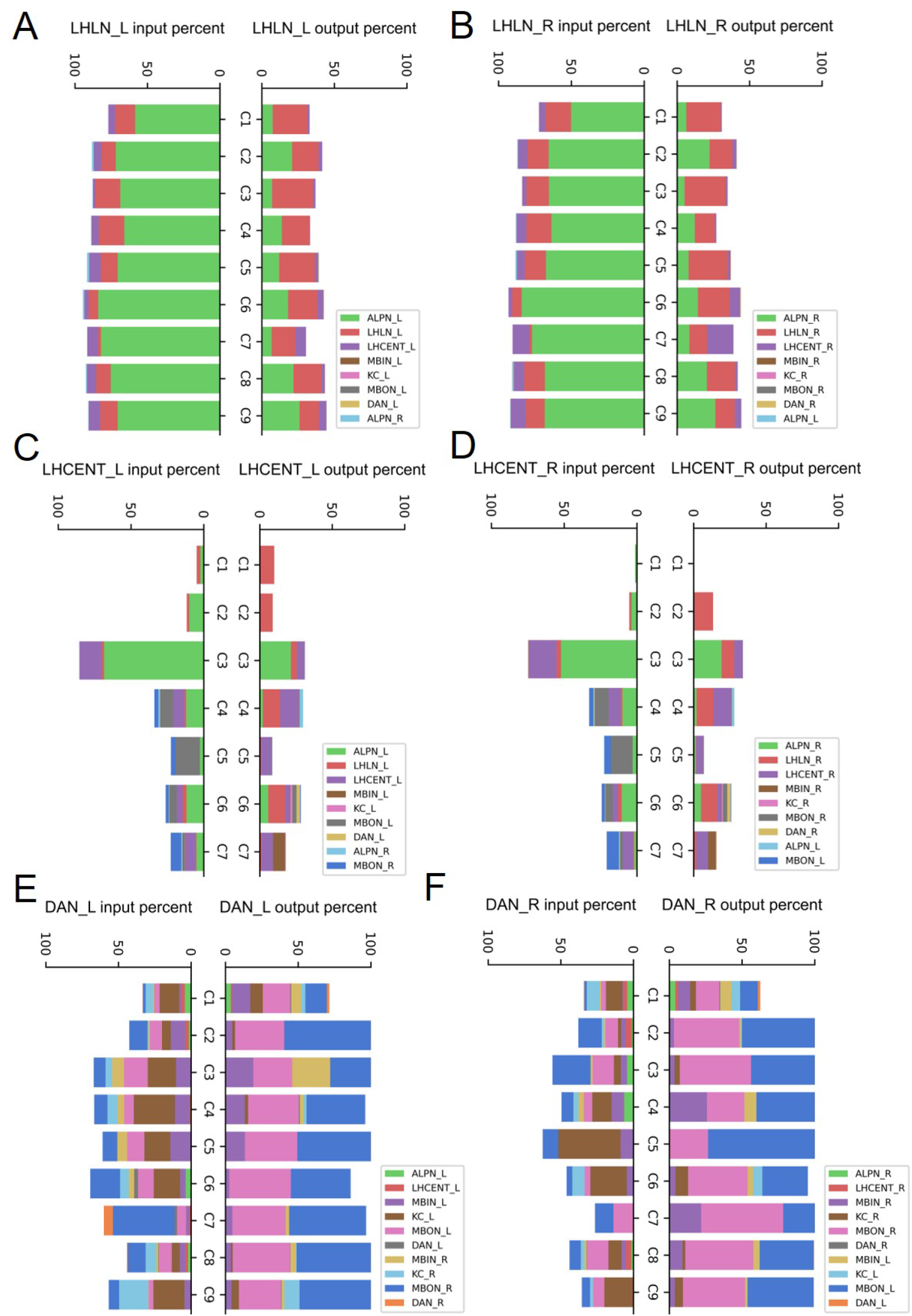
The percentage of synaptic input and output of clusters of the major neural classes. It considers the neural clusters within two hemispheres. (A, C, E) denote the results of the LHLNs, LHCENTs, and DANs in the left brain. (B, D, F) represent the results of these neural classes in the right brain.

**Supplementary Figure 9:**
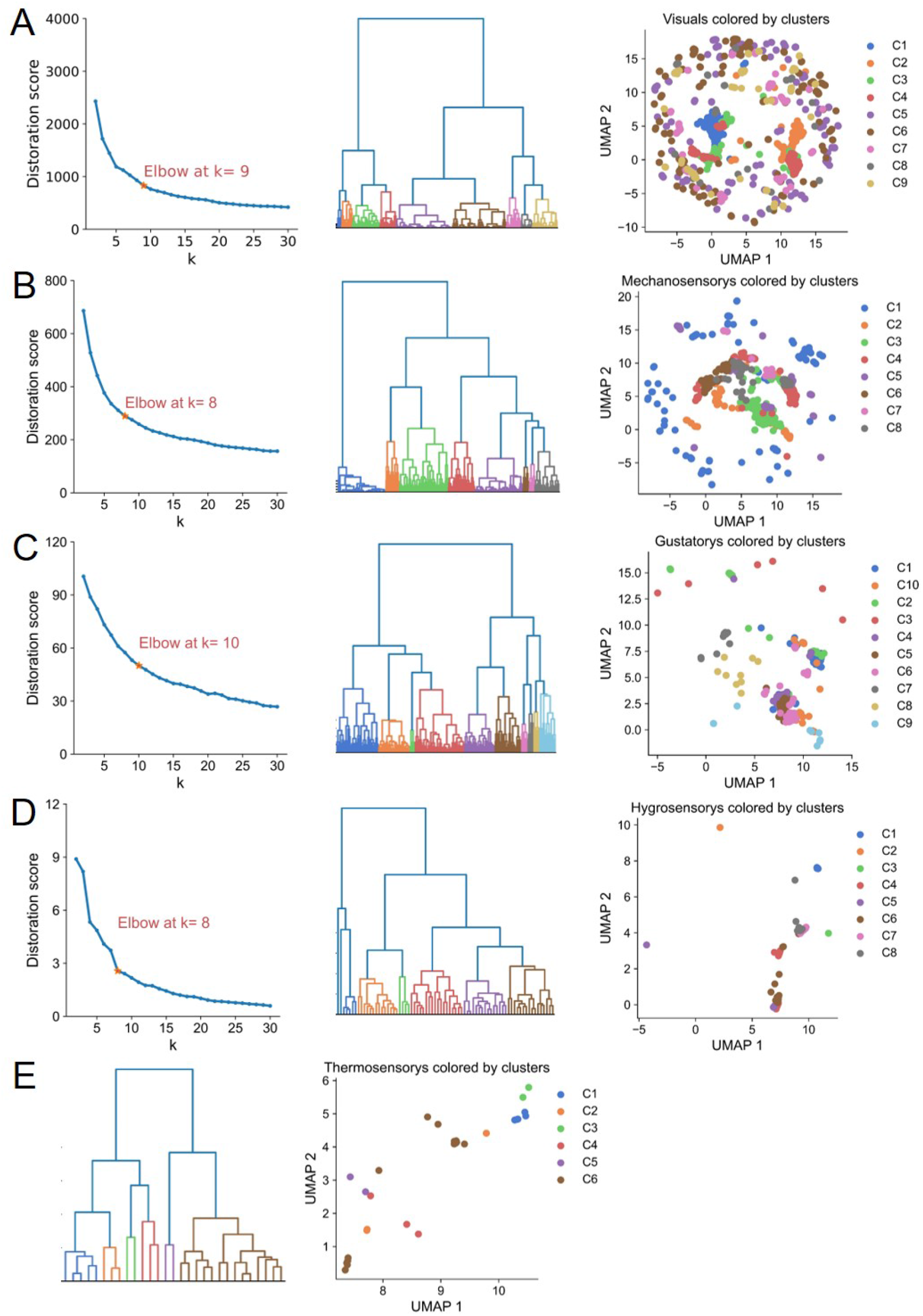
Hierarchical clustering results for the major sensory neurons. They include visual neurons (A), mechanosensory neurons (B), gustatory neurons (C), hygrosensory neurons (D) and thermosensory neurons (E).

**Supplementary Figure 10:**
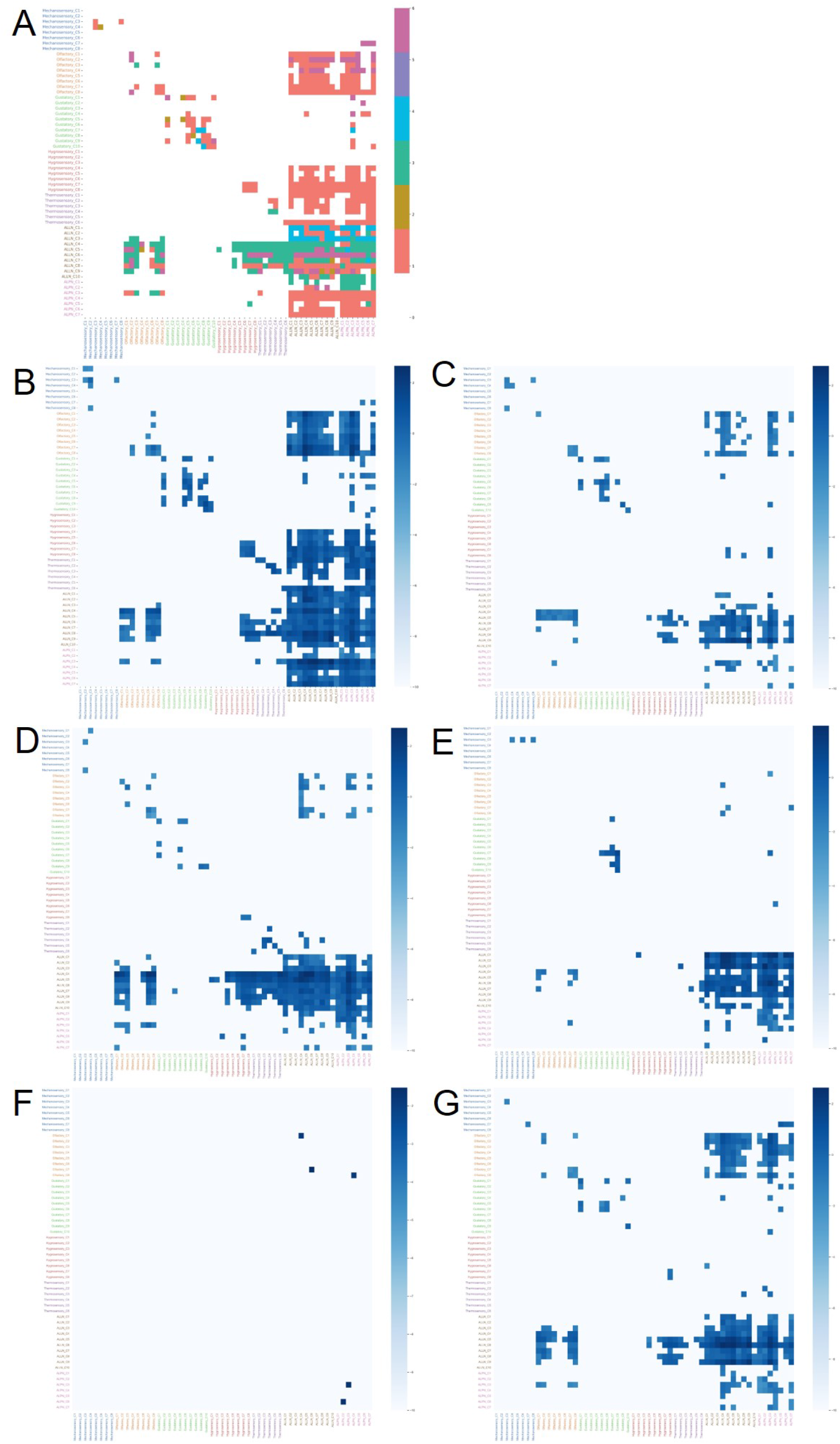
Connectivity matrix for neuronal clusters along the olfactory pathway. (A) The connectivity matrix of clusters for these sensory neuronal classes. Each element in the connectivity matrix represents the dominant synaptic type between two neural clusters. (B-G) The connectivity matrix of clusters solely for synapses with unique neurotransmitters, including ACH (B), DA (C), GABA (D), GLUT (E), OCT (F), SER (G).

## Notes

### Competing Interest Statement

The authors have declared no competing interest.

## References

[1] Hans Colonius and Adele Diederich. Formal models and quantitative measures of multisensory integration: a selective overview. European Journal of Neuroscience, 51(5):1161–1178, 2020.

[2] Timothy A Currier and Katherine I Nagel. Multisensory control of navigation in the fruit fly. Current Opinion in Neurobiology, 64:10–16, 2020.

[3] Mark T Wallace, Brian N Carriere, Thomas J Perrault, J William Vaughan, and Barry E Stein. The development of cortical multisensory integration. Journal of Neuroscience, 26(46):11844–11849, 2006.

[4] Greg SB Suh, Shlomo Ben-Tabou de Leon, Hiromu Tanimoto, André Fiala Seymour Benzer, and David J Anderson. Light activation of an innate olfactory avoidance response in drosophila. Current Biology, 17(10):905–908, 2007.

[5] Elizabeth C Marin, Laurin Büld, Maria Theiss, Tatevik Sarkissian, Ruairí JV Roberts, Robert Turnbull, Imaan FM Tamimi, Markus W Pleijzier, Willem J Laursen, Nik Drummond, et al. Connectomics analysis reveals first-, second-, and third-order thermosensory and hygrosensory neurons in the adult drosophila brain. Current Biology, 30(16):3167–3182, 2020.

[6] Jerome Carriot, Mohsen Jamali, and Kathleen E Cullen. Rapid adaptation of multisensory integration in vestibular pathways. Frontiers in systems neuroscience, 9:59, 2015.

[7] Manuel R Mercier and Celine Cappe. The interplay between multisensory integration and perceptual decision making. NeuroImage, 222:116970, 2020.

[8] Ilsong Choi, Ilayda Demir, Seungmi Oh, and Seung-Hee Lee. Multisensory integration in the mammalian brain: diversity and flexibility in health and disease. Philosophical Transactions of the Royal Society B, 378(1886):20220338, 2023.

[9] Ann-Shyn Chiang, Chih-Yung Lin, Chao-Chun Chuang, Hsiu-Ming Chang, Chang-Huain Hsieh, Chang-Wei Yeh, Chi-Tin Shih, Jian-Jheng Wu, Guo-Tzau Wang, Yung-Chang Chen, et al. Three-dimensional reconstruction of brain-wide wiring networks in drosophila at single-cell resolution. Current Biology, 21(1):1–11, 2011.

[10] Chi-Tin Shih, Olaf Sporns, Shou-Li Yuan, Ta-Shun Su, Yen-Jen Lin, Chao-Chun Chuang, Ting-Yuan Wang, Chung-Chuang Lo, Ralph J Greenspan, and Ann-Shyn Chiang. Connectomics-based analysis of information flow in the drosophila brain. Current Biology, 25(10):1249–1258, 2015.

[11] Zhihao Zheng, J Scott Lauritzen, Eric Perlman, Camenzind G Robinson, Matthew Nichols, Daniel Milkie, Omar Torrens, John Price, Corey B Fisher, Nadiya Sharifi, et al. A complete electron microscopy volume of the brain of adult drosophila melanogaster. Cell, 174(3):730–743, 2018.

[12] Louis K Scheffer, C Shan Xu, Michal Januszewski, Zhiyuan Lu, Shin-ya Takemura, Kenneth J Hayworth, Gary B Huang, Kazunori Shinomiya, Jeremy Maitlin-Shepard, Stuart Berg, et al. A connectome and analysis of the adult drosophila central brain. eLife, 9:e57443, 2020.

[13] Cory M Root, Julia L Semmelhack, Allan M Wong, Jorge Flores, and Jing W Wang. Propagation of olfactory information in drosophila. Proceedings of the National Academy of Sciences, 104(28):11826–11831, 2007.

[14] Yoichi Seki, Hany KM Dweck, Jürgen Rybak, Dieter Wicher, Silke Sachse, and Bill S Hansson. Olfactory coding from the periphery to higher brain centers in the drosophila brain. BMC Biology, 15:1–20, 2017.

[15] Shahar Frechter, Alexander Shakeel Bates, Sina Tootoonian, Michael-John Dolan, James Manton, Arian Rokkum Jamasb, Johannes Kohl, Davi Bock, and Gregory Jefferis. Functional and anatomical specificity in a higher olfactory centre. eLife, 8:e44590, 2019.

[16] Quentin Gaudry, Elizabeth J Hong, Jamey Kain, Benjamin L de Bivort, and Rachel I Wilson. Asymmetric neurotransmitter release enables rapid odour lateralization in drosophila. Nature, 493(7432):424–428, 2013.

[17] Sonia G Chin, Sarah E Maguire, Paavo Huoviala, Gregory SXE Jefferis, and Christopher J Potter. Olfactory neurons and brain centers directing oviposition decisions in drosophila. Cell Reports, 24(6):1667–1678, 2018.

[18] Vanessa Ruta, Sandeep Robert Datta, Maria Luisa Vasconcelos, Jessica Freeland, Loren L Looger, and Richard Axel. A dimorphic pheromone circuit in drosophila from sensory input to descending output. Nature, 468(7324):686–690, 2010.

[19] E Josephine Clowney, Shinya Iguchi, Jennifer J Bussell, Elias Scheer, and Vanessa Ruta. Multimodal chemosensory circuits controlling male courtship in drosophila. Neuron, 87(5):1036–1049, 2015.

[20] Sven Dorkenwald, Peter H Li, Michał Januszewski Daniel R Berger, Jeremy Maitin-Shepard, Agnes L Bodor, Forrest Collman, Casey M Schneider-Mizell, Nuno Maçarico da Costa, Jeff W Lichtman, et al. Multi-layered maps of neuropil with segmentation-guided contrastive learning. Nature Methods, pages 1–10, 2023.

[21] Michael Winding, Benjamin D Pedigo, Christopher L Barnes, Heather G Patsolic, Youngser Park, Tom Kazimiers, Akira Fushiki, Ingrid V Andrade, Avinash Khandelwal, Javier Valdes-Aleman, et al. The connectome of an insect brain. Science, 379(6636):eadd9330, 2023.

[22] Marta Costa, James D Manton, Aaron D Ostrovsky, Steffen Prohaska, and Gregory SXE Jefferis. Nblast: rapid, sensitive comparison of neuronal structure and construction of neuron family databases. Neuron, 91(2):293–311, 2016.

[23] Jacob C Worrell, Jeffrey Rumschlag, Richard F Betzel, Olaf Sporns, and Bratislav Mišić. Optimized connectome architecture for sensory-motor integration. Network Neuroscience, 1(4):415–430, 2017.

[24] Rafael Yuste, Rosa Cossart, and Emre Yaksi. Neuronal ensembles: Building blocks of neural circuits. Neuron, 2024.

[25] Dan Busbridge, Dane Sherburn, Pietro Cavallo, and Nils Y. Hammerla. Relational graph attention networks, 2019.

[26] Marc Brockschmidt. Gnn-film: Graph neural networks with feature-wise linear modulation. In International Conference on Machine Learning, pages 1144–1152. PMLR, 2020.

[27] Filippo Maria Bianchi. Simplifying clustering with graph neural networks. Proceedings of the Northern Lights Deep Learning Workshop 2023, 2023.

[28] Michael Schlichtkrull, Thomas N Kipf, Peter Bloem, Rianne van den Berg, Ivan Titov, and Max Welling. Modeling relational data with graph convolutional networks. In European Semantic Web Conference, pages 593–607. Springer, 2018.

[29] Shichao Zhu, Chuan Zhou, Shirui Pan, Xingquan Zhu, and Bin Wang. Relation structure-aware heterogeneous graph neural network. In 2019 IEEE international conference on data mining (ICDM), pages 1534–1539. IEEE, 2019.

[30] Thomas N. Kipf and Max Welling. Semi-supervised classification with graph convolutional networks. In International Conference on Learning Representations, 2017.

[31] Petar Veličković, Guillem Cucurull, Arantxa Casanova, Adriana Romero, Pietro Liò, and Yoshua Bengio. Graph attention networks. In International Conference on Learning Representations, 2018.

[32] Will Hamilton, Zhitao Ying, and Jure Leskovec. Inductive representation learning on large graphs. Advances in neural information processing systems, 30, 2017.

[33] Zonghan Wu, Shirui Pan, Fengwen Chen, Guodong Long, Chengqi Zhang, and S Yu Philip. A comprehensive survey on graph neural networks. IEEE transactions on neural networks and learning systems, 32(1):4–24, 2020.

[34] Paul Y Wang, Sandalika Sapra, Vivek Kurien George, and Gabriel A Silva. Generalizable machine learning in neuroscience using graph neural networks. Frontiers in artificial intelligence, 4:618372, 2021.

[35] Alaa Bessadok, Mohamed Ali Mahjoub, and Islem Rekik. Graph neural networks in network neuroscience. IEEE Transactions on Pattern Analysis and Machine Intelligence, 45(5):5833–5848, 2022.

[36] Gabriele Corso, Hannes Stark, Stefanie Jegelka, Tommi Jaakkola, and Regina Barzilay. Graph neural networks. Nature Reviews Methods Primers, 4(1):17, 2024.

[37] Xiyang Sun and Fumiyasu Komaki. Bhgnn-rt: Network embedding for directed heterogeneous graphs. arXiv preprint 2311.14404, 2023.

[38] Jacob Tanner, Joshua Faskowitz, Andreia Sofia Teixeira, Caio Seguin, Ludovico Coletta, Alessandro Gozzi, Bratislav Mišić, and Richard F Betzel. A multi-modal, asymmetric, weighted, and signed description of anatomical connectivity. Nature Communications, 15(1):5865, 2024.

[39] Albert Lin, Runzhe Yang, Sven Dorkenwald, Arie Matsliah, Amy R Sterling, Philipp Schlegel, Szi-chieh Yu, Claire E McKellar, Marta Costa, Katharina Eichler, et al. Network statistics of the whole-brain connectome of drosophila. Nature, 634(8032):153–165, 2024.

[40] Petar Velickovic, William Fedus, William L Hamilton, Pietro Liò, Yoshua Bengio, and R Devon Hjelm. Deep graph infomax. ICLR (Poster), 2(3):4, 2019.

[41] Toshihide Hige, Yoshinori Aso, Mehrab N Modi, Gerald M Rubin, and Glenn C Turner. Heterosynaptic plasticity underlies aversive olfactory learning in drosophila. Neuron, 88(5):985–998, 2015.

[42] Shin-ya Takemura, Yoshinori Aso, Toshihide Hige, Allan Wong, Zhiyuan Lu, C Shan Xu, Patricia K Rivlin, Harald Hess, Ting Zhao, Toufiq Parag, et al. A connectome of a learning and memory center in the adult drosophila brain. eLife, 6:e26975, 2017.

[43] Suewei Lin. The making of the drosophila mushroom body. Frontiers in Physiology, 14:1091248, 2023.

[44] Louis K Scheffer. Evidence of wiring development processes from the connectome of adult drosophila. bioRxiv, pages 2021–05, 2021.

[45] Philipp Schlegel, Yijie Yin, Alexander S Bates, Sven Dorkenwald, Katharina Eichler, Paul Brooks, Daniel S Han, Marina Gkantia, Marcia Dos Santos, Eva J Munnelly, et al. Whole-brain annotation and multi-connectome cell typing of drosophila. Nature, 634(8032):139–152, 2024.

[46] Yoshinori Aso, Daisuke Hattori, Yang Yu, Rebecca M Johnston, Nirmala A Iyer, Teri-TB Ngo, Heather Dionne, LF Abbott, Richard Axel, Hiromu Tanimoto, et al. The neuronal architecture of the mushroom body provides a logic for associative learning. eLife, 3:e04577, 2014.

[47] Yoshinori Aso, Divya Sitaraman, Toshiharu Ichinose, Karla R Kaun, Katrin Vogt, Ghislain Belliart-Guérin, Pierre-Yves Plaçais, Alice A Robie, Nobuhiro Yamagata, Christopher Schnaitmann, et al. Mushroom body output neurons encode valence and guide memory-based action selection in drosophila. eLife, 3:e04580, 2014.

[48] Feng Li, Jack W Lindsey, Elizabeth C Marin, Nils Otto, Marisa Dreher, Georgia Dempsey, Ildiko Stark, Alexander S Bates, Markus William Pleijzier, Philipp Schlegel, et al. The connectome of the adult drosophila mushroom body provides insights into function. eLife, 9:e62576, 2020.

[49] Ketan Mehta, Rebecca F Goldin, and Giorgio A Ascoli. Circuit analysis of the drosophila brain using connectivity-based neuronal classification reveals organization of key communication pathways. Network Neuroscience, 7(1):269–298, 2023.

[50] Hiroaki Kitano. Biological robustness. Nature Reviews Genetics, 5(11):826–837, 2004.

[51] Gabor Timár, Alexander V Goltsev, Sergey N Dorogovtsev, and José FF Mendes. Mapping the structure of directed networks: Beyond the bow-tie diagram. Physical Review Letters, 118(7):078301, 2017.

[52] Peter Csermely, András London, Ling-Yun Wu, and Brian Uzzi. Structure and dynamics of core/periphery networks. Journal of Complex Networks, 1(2):93–123, 2013.

[53] François Lapraz, Cloé Fixary-Schuster, and Stéphane Noselli. Brain bilateral asymmetry–insights from nematodes, zebrafish, and drosophila. Trends in Neurosciences, 2024.

[54] Rashmit Kaur, Michael Surala, Sebastian Hoger, Nicole Grössmann, Alexandra Grimm, Lorin Timaeus, Wolfgang Kallina, and Thomas Hummel. Pioneer interneurons instruct bilaterality in the drosophila olfactory sensory map. Science Advances, 5(10):eaaw5537, 2019.

[55] Sven Dorkenwald, Arie Matsliah, Amy R Sterling, Philipp Schlegel, Szi-Chieh Yu, Claire E McKellar, Albert Lin, Marta Costa, Katharina Eichler, Yijie Yin, et al. Neuronal wiring diagram of an adult brain. Nature, 634(8032):124–138, 2024.

[56] Andrew Kachites McCallum, Kamal Nigam, Jason Rennie, and Kristie Seymore. Automating the construction of internet portals with machine learning. Information Retrieval, 3:127–163, 2000.

[57] C Lee Giles, Kurt D Bollacker, and Steve Lawrence. Citeseer: An automatic citation indexing system. In Proceedings of the third ACM conference on Digital libraries, pages 89–98, 1998.

[58] Julian McAuley, Christopher Targett, Qinfeng Shi, and Anton Van Den Hengel. Image-based recommendations on styles and substitutes. In Proceedings of the 38th international ACM SIGIR conference on research and development in information retrieval, pages 43–52, 2015.

[59] Aaron Clauset, Cosma Rohilla Shalizi, and Mark EJ Newman. Power-law distributions in empirical data. SIAM Review, 51(4):661–703, 2009.

[60] Michele Migliore and Gordon M Shepherd. An integrated approach to classifying neuronal phenotypes. Nature Reviews Neuroscience, 6(10):810–818, 2005.

[61] D Anderson and K Burnham. Model selection and multi-model inference. Second. NY: Springer-Verlag, 63(2020):10, 2004.

[62] Jianxi Luo and Daniel E Whitney. Asymmetry in in-degree and out-degree distributions of large-scale industrial networks. Structure and Dynamics, 8(2), 2015.

[63] Natsuhiro Ichinose, Tetsushi Yada, and Hiroshi Wada. Asymmetry in indegree and outdegree distributions of gene regulatory networks arising from dynamical robustness. Physical Review E, 97(6):062315, 2018.

[64] Travis A Jarrell, Yi Wang, Adam E Bloniarz, Christopher A Brittin, Meng Xu, J Nichol Thomson, Donna G Albertson, David H Hall, and Scott W Emmons. The connectome of a decision-making neural network. science, 337(6093):437–444, 2012.

[65] Sen Song, Per Jesper Sjöström, Markus Reigl, Sacha Nelson, and Dmitri B Chklovskii. Highly nonrandom features of synaptic connectivity in local cortical circuits. PLoS Biology, 3(3):e68, 2005.

[66] Nicolas X Tritsch, Adam J Granger, and Bernardo L Sabatini. Mechanisms and functions of gaba co-release. Nature Reviews Neuroscience, 17(3):139–145, 2016.

[67] Nils Eckstein, Alexander Shakeel Bates, Andrew Champion, Michelle Du, Yijie Yin, Philipp Schlegel, Alicia Kun-Yang Lu, Thomson Rymer, Samantha Finley-May, Tyler Paterson, et al. Neurotransmitter classification from electron microscopy images at synaptic sites in drosophila melanogaster. Cell, 187(10):2574–2594, 2024.

[68] Chun Wang, Shirui Pan, Ruiqi Hu, Guodong Long, Jing Jiang, and Chengqi Zhang. Attributed graph clustering: a deep attentional embedding approach. In Proceedings of the 28th International Joint Conference on Artificial Intelligence, pages 3670–3676, 2019.

[69] Costas Mavromatis and George Karypis. Graph infoclust: Maximizing coarse-grain mutual information in graphs. In Pacific-Asia Conference on Knowledge Discovery and Data Mining, pages 541–553. Springer, 2021.

